# Quantitative trait locus mapping of osmotic stress response in the fungal wheat pathogen *Zymoseptoria tritici*

**DOI:** 10.1101/2022.06.07.495078

**Authors:** Jessica Stapley, Bruce A. McDonald

## Abstract

Osmotic stress is a ubiquitous and potent stress for all living organisms, but few studies have investigated the genetic basis of salt tolerance in filamentous fungi. The main aim of this study was to identify regions of the genome associated with tolerance to potassium chloride in the wheat pathogen *Zymoseptoria tritici.* A secondary aim was to identify candidate genes affecting salt tolerance within the most promising chromosomal regions. We achieved these aims with a quantitative trait locus (QTL) mapping study using offspring from two crosses grown in vitro in the presence or absence of osmotic stress imposed by 0.75M KCl. We identified significant QTL for most of the traits in both crosses. Several QTL overlapped with QTL identified in earlier studies for other traits and some QTL explained trait variation in both the control and salt stress environments. A significant QTL on chromosome 3 explained variation in colony radius at 8 days post inoculation (dpi) in the KCl environment as well as colony radius KCl tolerance at 8 dpi. The QTL peak had a high LOD and encompassed an interval containing only 36 genes. Six of these genes present promising candidates for functional analyses. A GO enrichment analysis of QTL unique to the KCl environment found evidence for enrichment of functions involved in osmotic stress responses.

## Introduction

Osmotic stress is a common and potent environmental stress that can limit growth and reproduction in many organisms. At the cellular level osmotic stress results in changes to the movement of water into or out of the cell through osmosis. To counteract osmotic stress, organisms need to accumulate compatible solutes (e.g. glycerol, erythritol, arabitol, mannitol and trehalose) – that can balance osmosis and provide turgor pressure (Kogej *et al*. 2007; Araújo *et al*. 2020). In fungal pathogens of plants, the pathogen may be exposed to a relatively high salt concentration (0.5M) within the plant (see Araújo et al., 2020) as well as experience changes in salt and osmolyte concentrations as plant tissues degrade (Jacob *et al*. 2015). The genes and pathways involved in responding to the resulting osmotic stresses may also affect plant-pathogen interactions and virulence. For example, turgor pressure which is mediated by changes in vacuole size and number is important in both plant tissue penetration and response to osmotic stress (Richards *et al*. 2010). Genes involved in cell wall integrity and remodelling, e.g. glycoside hydrolases, can mediate osmotic stress response and virulence (Li *et al*. 2016). Voltage-gated chloride channels, which are involved in the regulation of cell volume, also influence virulence in fungal pathogens of both plants and humans (Zhu and Williamson 2003; Cañero and Roncero 2008). Knowledge of the genetic basis of osmotic sensitivity in plant pathogens can thus provide insights into the genetic basis of environmental stress responses as well as disease development.

The immediate effects of osmotic stress in fungi include stalled growth, changes in the recruitment of water, collapse of the cytoskeleton, altered plasma membranes and a reduction in glycerol transport (Hohmann 2002; Duran *et al*. 2010). Longer-term adaptation to osmotic stress involves modifications of metabolism, plasma membranes, cell wall thickness and cell wall melanisation, as well as an accumulation of mycosporines (Hohmann 2002; Kogej *et al*. 2007). Osmotic stress is also likely to activate many genes and pathways involved in the ‘general stress response’. For example, up to 10% of the genes that were differentially expressed under different individual stressors were shared across different osmotic stress treatments (Hohmann 2002). Stress response regulators and genes such as HSP90 and GTPase *Ras1* are considered promising antifungal drug targets for human pathogens (LeBlanc *et al*. 2020). These proteins may also become fungicide targets for plant pathogens as our understanding of the molecular mechanisms governing stress responses in plant pathogenic fungi improves.

Although the key salt stress response pathways and genes have been identified and well characterized in model yeasts, studies in filamentous fungi could provide novel insights into the genes involved in responding to osmotic stress. The HOG pathway is an osmoresponsive system and one of the most well-understood mitogen-activated protein (MAP) kinase pathways (Hohmann 2002). In yeast, two genes are central components of this pathway – Pbs2 and Hog1p kinase. In plant pathogens, HOG signalling appears more complex than in yeast, although it is still poorly understood (Jacob *et al*. 2015). Several filamentous pathogenic fungi, including our model *Zymoseptoria tritici*, have multiple (up to 17) histidine kinases, whereas *S. cerevisiae* has only one (Sln1p) (Jacob *et al*. 2015). Several studies have identified transcription factors (TFs) involved in regulating the fungal response to salt stress, as well as other stressors - including temperature and fungicide stress. For example, in *Aspergillus oryzae*, GATA TFs were shown to be important in responding to salt and temperature stress (Jiang *et al*. 2021). In the causal agent of Fusarium head blight (*Fusarium graminearum*), the nutrient and stress factor 1 (NSf1) C_2_H_2_ zinc fingers protein plays an important role in salt stress response, sexual-asexual reproduction, vegetative growth, fungicide sensitivity and pathogenicity (Shi *et al*. 2021), and a Zn(2)-C6 fungal-type DNA-binding TF (Fss1) is required for sodium and lithium tolerance (Son *et al*. 2015). Cell wall melanisation also appears to confer benefits for osmotic stress tolerance. For example, cell wall melanisation in halotolerant fungi helps to maintain high concentrations of glycerol and contain the resulting turgor pressure by strengthening and preventing leakage through the cell wall (Kogej *et al*. 2007). These examples coming from a relatively small number of fungal species demonstrate how the molecular mechanisms of salt response can vary across taxa and illustrate that additional work is needed to understand how filamentous fungi adapt to salt stress.

To improve our understanding of osmotic stress response in fungi we performed a quantitative trait loci (QTL) mapping study using the wheat pathogen *Z. tritici* growing under salt stress. QTL studies have been instrumental in finding candidate genes for multiple traits in this widespread and damaging wheat pathogen, including melanisation (Lendenmann *et al*. 2014), fungicide sensitivity (Lendenmann *et al*. 2015), morphological switching (Francisco *et al*. 2023) and oxidative stress (Zhong *et al*. 2021), and enabled the cloning and functional validation of genes affecting virulence (Zhong et al., 2017; Meile et al. 2018), melanisation (Krishnan et al. 2018), and morphological switching (Francisco *et al*. 2023). QTL mapping identifies genomic regions that explain variation in a trait and that are likely to harbour genes responsible for creating variation in the trait. The aim of this study was to identify large effect loci associated with tolerance to salt stress in *Z. tritici* and identify plausible candidate genes within the most promising QTL intervals.

## Methods

### QTL mapping crosses

Parent and progeny isolates used in this study are from two crosses that were described previously (Lendenmann *et al*. 2014). The original choice of strains for making crosses was based upon a comprehensive phenotyping campaign of an international collection of 150 strains, including 30 strains from Switzerland (Zhan *et al*. 2005). In this original phenotyping campaign, we measured seven traits for each strain, including in planta virulence and pycnidia size and density, and growth rates in vitro under different temperatures and in the presence of fungicides. Based on data from these seven traits, four Swiss strains were selected to make the QTL mapping crosses that yielded 700 offspring over the two crosses. The four parents have since been tested under other environmental stressors (e.g. reactive oxygen (Zhong *et al*. 2021)) including KCl (0.25-1.0M). In the presence of 0.75M KCl the parents differed in both growth and melanisation traits (unpublished data). The crosses were made by coinfecting wheat leaves with the parent strains, with the resulting ascospores collected from infected leaves and grown in vitro. One cross was between ST99CH1A5 and ST99CH1E4 (herein referred to as 1A5 and 1E4 respectively) and the other was between ST99CH3D7 and ST99CH3D1 (herein referred to as 3D7 and 3D1 respectively).

### Genotyping

SNP genotype data for the progeny were obtained from a RAD sequence dataset that was previously produced in our lab (first described in (Lendenmann *et al*. 2014). In brief, the genome was cut using the restriction enzyme *Pst*l and the libraries were sequenced on an Illumina HiSeq2000 with paired-end sequencing. Complete genome sequences of the parental strains (Croll *et al*. 2013)(NCBI Biosample: SRS383146 (ST99CH3D1), SRS383147 (ST99CH3D7), SRS383142 (ST99CH1A5), and SRS383143 (ST99CH1E4)) were used to SNP genotype the parents and offspring.

RADseq processing and variant discovery is described in detail at https://github.com/jessstapley/QTL-mapping-Z.-tritici. In brief, RADseq reads were trimmed of adaptors and low-quality sequence using trimmomatic (v0.35). The RADseq reads were mapped to a reference genome using bwa mem (v0.7.17). Reads from progeny of the 3D7×3D1 cross were mapped to the 3D7 reference genome and progeny from the 1A5×1E4 cross were mapped to the 1A5 reference genome. Variant calling was done using the GATK Germline Short Variant Discovery pipeline, following their Best Practice recommendations (https://gatk.broadinstitute.org/hc/en-us/sections/360007226651-Best-Practices-Workflows). After this we applied the following filters: only biallelic SNPs, parents had alternative alleles, <50% missing genotypes per marker, <50% missing genotypes per individual, mean read depth >3 and <30, depth quality (QD)>5, mapping quality >40, minor allele frequency >0.02 and <0.80, and SNPs in regions where the SNP density is >3 in 10bp were removed.

### Generating linkage maps

Linkage maps were made using R (R Core Team 2020), R Studio (RStudio Team 2020) and the R package qtl (v1.48-1 (Broman *et al*. 2003)). As the order and base pair positions of the markers were known from the reference genomes, we only needed to calculate the recombination distance between markers to create the linkage map. Before map construction, we removed putative clones. To do this we calculated genetic similarity using the ‘comparegeno’ function and removed individuals with >99% shared genotypes (83 progeny were removed from the 1A5×1E4 cross, 74 progeny were removed from the 3D7×3D1 cross). The linkage map was constructed for each chromosome separately using the ‘estimatemap’ function with method ‘morgan’. Then we performed multiple rounds of cleaning to remove error-prone markers that caused map inflation. First, we removed markers that had an estimated recombination distance above the 99.9% quantile. Then recombination rate was estimated again and we inspected the maps. In regions of the map where there was evidence of map inflation and spurious double recombination events, we visualised these regions in an Integrated Genome Viewer (IGV, https://igv.org) to look at the mapping alignments. Markers were removed from regions with poor alignments and evidence of mismapping. A summary of the two linkage maps is provided in Supplementary Table S1 and the complete maps are available at (https://github.com/jessstapley/QTL-mapping-Z.-tritici). After applying all of these filters, the average marker spacing was 1217 bp (0.19 cM) and 529 bp (0.14 cM) for the 1A5×1E4 and 3D7×3D1 crosses, respectively.

### Phenotyping

We compared colony growth and melanisation in a control environment and in the presence of potassium chloride (KCl) to simulate a salt stress environment. Potassium chloride (KCl) has been used extensively in the study of stress responses in fungi because it is environmentally ubiquitous and lacks the toxic effects of Na^+^ (Hohmann 2002; Araújo *et al*. 2020). Only non-clonal offspring strains were phenotyped (for 1A5×1E4 n= 259 strains; for 3D7×3D1 n=265 strains). The basic protocols for isolate recovery from -80^0^C storage, growth in vitro and the measurements of colony size and colony melanisation were described previously (Zhong *et al*. 2021). In brief, Petri dishes containing Difco potato dextrose agar (PDA) were inoculated with 200μl of a spore solution (concentration 200 spores/ml) and grown at 18^0^C for 8 and 12 days. In the salt stress environment, a sterile potassium chloride solution was added to the cooled and sterilized PDA to make a final concentration of 0.75M. Three replicates were grown in both the control and salt (KCl) environments.

Two traits were measured; colony area and colony grey value, using automated image analysis as described previously (Lendenmann *et al*. 2014; Zhong *et al*. 2021). Grey value was measured on a scale of 0-255, where darker, more melanised colonies have lower values (0 = black, 255 = white). Mean colony area (mm^2^) and grey value were calculated from measurements of multiple colonies on each plate and then the mean was calculated across the three replicate plates to obtain a final mean colony area and mean grey value for each strain in the two environments. Colony area was converted to colony radius by dividing by π and taking the square root of this value. Measurements were taken at 8- and 12-days post inoculation (dpi). For the analysis we used three different types of data; (1) time point/age: the colony radius and grey value at 8 and 12 dpi; (2) daily rate measurements: the change in colony radius or grey value between 8 and 12 dpi (i.e. the difference in colony radius between 8 and 12 dpi/difference in days); and (3) KCl tolerance measurements: the ratio of a measurement (colony radius, grey value, growth rate or melanisation rate) between the KCl environment and the control environment (i.e. colony radius KCl tolerance at 8 dpi = colony radius at 8 dpi in the KCl environment/colony radius at 8 dpi in the control environment, or growth rate KCl tolerance = growth rate in the KCl environment/growth rate in the control environment). For the KCl tolerance measurements, values >1 indicate that for that trait an isolate is more tolerant to KCl (higher value under salt stress), and values <1 indicate that for that trait measured an isolate is more sensitive to KCl. In total there were 18 traits (Table 1).

**Table 1.** Summary of the QTL mapping results for all traits. Grey value and colony radius were measured at two time points: 8 and 12 dpi. Growth rate was calculated as the change in colony radius/number of days. Melanisation rate was calculated as the change in grey value/number of days. Colony radius/grey value KCl tolerance were calculated by dividing the value (grey value or colony radius) in the control environment by the value in the KCl environment. Growth rate/Melanisation KCl tolerance were calculated by dividing the rate (melanisation or growth) in the control environment by the rate in the KCl environment. The QTL peak columns show the chromosomal position of significant QTL peak for each trait in the 3D7×3D1 and 1A5×1E4 crosses respectively, and the final column contains estimated heritability (h^2^) for each trait.

| Data Type | Environment | DPI | Trait | Trait Full Name | QTL peak<br>3D7x3D1 | $h^2$ | QTL peak<br>1A5x1E4 | $h^2$ |
| --- | --- | --- | --- | --- | --- | --- | --- | --- |
| Time point | Control | 8 | Colony radius | Colony radius in Control at 8dpi | 3,8,10,11 | 0.53 | 2,3,8 | 0.56 |
| Time point | Control | 12 | Colony radius | Colony radius in Control at 12dpi | 3,7,8,10,11 | 0.62 | 7,8 | 0.48 |
| Time point | KCl | 8 | Colony radius | Colony radius in KCl at 8dpi | 3,4 | 0.64 | 6, | 0.50 |
| Time point | KCl | 12 | Colony radius | Colony radius in KCl at 12dpi | 3,8 | 0.60 | 1,8 | 0.52 |
| Time point | Control | 8 | Grey value | Grey value in Control at 8dpi | 8,10,11 | 0.42 | 1,2,3,8 | 0.63 |
| Time point | Control | 12 | Grey value | Grey value in Control at 12dpi | 11 | 0.22 | 2,3,8 | 0.44 |
| Time point | KCl | 8 | Grey value | Grey value in KCl at 8dpi | 8,11 | 0.54 | 8 | 0.34 |
| Time point | KCl | 12 | Grey value | Grey value in KCl at 12dpi | 1,2,11 | 0.34 | No Sig. QTL | 0.29 |
| Rate | Control | 8-12 | Growth rate | Growth rate in Control | 3,8,11 | 0.61 | 7,8 | 0.33 |
| Rate | KCl | 8-12 | Growth rate | Growth rate in KCl | 3,8,11 | 0.45 | 8, | 0.49 |
| Rate | Control | 8-12 | Melanisation rate | Melanisation rate in Control | 10 | 0.28 | 2,3,12 | 0.54 |
| Rate | KCl | 8-12 | Melanisation rate | Melanisation rate in KCl | 1,6 | 0.32 | No Sig. QTL | 0.27 |
| Tolerance | Both | 8 | Grey value | Grey value KCl tolerance at 8 dpi | 8,10 | 0.37 | 1,2,3,8 | 0.58 |
| Tolerance | Both | 12 | Grey value | Grey value KCl tolerance at 12 dpi | 11 | 0.18 | No Sig. QTL | 0.34 |
| Tolerance | Both | 8 | Colony radius | Colony radius KCl tolerance at 8dpi | 3,4,10 | 0.59 | 8 | 0.42 |
| Tolerance | Both | 12 | Colony radius | Colony radius KCl tolerance at 12dpi | 3,10,11 | 0.47 | 1 | 0.37 |
| Tolerance | Both | 8-12 | Growth rate | Growth rate KCl tolerance | 11 | 0.40 | 1,5 | 0.45 |
| Tolerance | Both | 8-12 | Melanisation rate | Melanisation rate KCl tolerance | 1 | 0.14 | No Sig. QTL | 0.32 |

### QTL mapping and analysis of genes within QTL intervals

QTL mapping was performed using the R (v 3.6.0) package ‘qtl2’ (v2_0.24) as described in detail on github (https://github.com/jessstapley/QTL-mapping-Z.-tritici). We scanned the genome for a single QTL per chromosome with the ‘*scan1*’ function using a linear mixed effect model, and we controlled for the relatedness of individuals (i.e. we included a random polygenic effect) by fitting the kinship matrix as a random effect. Models that take into account the genetic covariance between individuals can reduce the false discovery rate in QTL scans and outperform models that do not include this information in the model (Malosetti *et al*. 2011). The significance threshold for a QTL peak was determined by permutation tests (n=1000) and we calculated a Bayes Credible Interval (95%) to identify the interval size around the QTL peak. We used custom R scripts to extract a list of the genes within the interval from the annotation (gff) files of each reference genome. The putative encoded function of each gene was determined in previous analyses ((for 1A5 see (Plissonneau *et al*. 2018), for 3D7 see (Plissonneau *et al*. 2016)). The annotations were obtained using Interproscan (https://www.ebi.ac.uk/interpro/) against multiple protein databases and then screened using multiple methods (e.g. Signal https://services.healthtech.dtu.dk/service.php?SignalP-5.0, Phobius https://phobius.sbc.su.se/) to determine if the encoded proteins contained likely signal peptides or secreted domains. The putative effect of a SNP in the coding sequence was determined using SNPeff (http://pcingola.github.io/SnpEff/). Pathway (GO and KEGG) enrichment analysis was performed for each trait with a significant QTL that was unique to the KCl environment following a tutorial for non-model species (https://archetypalecology.wordpress.com/2021/01/27/how-to-perform-kegg-and-go-enrichment-analysis-of-non-model-species-using-r/). GO annotations were retrieved from the annotation files describe above. For GO enrichment the R package ‘topGo’ (Alexa and Rahnenfuhrer 2023) was used to perform a Fischer test on the genes within each QTL interval following the guidelines in the vignette. All three ontologies – Biological Process (BP), Cellular Component (CC) and Molecular Function (MF) were analysed. For KEGG pathways enrichment, GhostKOALA (https://www.kegg.jp/ghostkoala/) was used to find K numbers for all transcripts. The program found K numbers for 34% (4091/11726) and 49% (5970/12072) of transcripts for 3D7 and 1A5 respectively. The enrichment test was performed using the R package clusterProfiler (Wu *et al*. 2021) and the ‘enricher’ function. The background set for both GO and KEGG was the entire gene/transcript set for each reference genome (3D7 and 1A5).

### Identifying putative candidate genes in the 3D7×3D1 cross

Genes within a single QTL interval on chromosome 3 were further investigated to identify the most likely candidate genes responsible for the phenotype. We identified the orthologous genes in the 3D1 parent using an analysis performed across the genomes of 19 reference strains (Badet *et al*. 2020). We inspected the alignments in the region of interest in IGV. We also blasted the 3D1 gene sequences (including ±200bp flanking regions) against the 3D7 genome to calculate sequence similarity and identify indels.

### RNAseq data analysis

We analysed RNAseq data that was previously created for the four parental strains during infection of the susceptible wheat cultivar Drifter (Palma-Guerrero *et al*. 2017) as well as cultures grown in vitro on two types of liquid media (yeast sucrose broth (YSB): yeast extract 10 g/L and sucrose 10 g/L, pH 6.8; and carbon-depleted minimal medium (MM), pH 5.8 (Francisco *et al*. 2019)). The raw sequence reads were downloaded from the Short Read Archive (SRA, https://www.ncbi.nlm.nih.gov/sra, Bioproject: *invitro* SRP152081, *invivo* SRP077418). RNAseq data processing and analysis is described in detail at github (https://github.com/jessstapley/QTL-mapping-Z.-tritici). In brief, reads were trimmed and then mapped to one of two reference genomes (reads from 3D1 and 3D7 were mapped to the 3D7 reference genome while reads from 1E4 and 1A5 were mapped to the 1A5 reference genome). Then the number of reads mapping to each gene was counted using R::Rsubread (Liao *et al*. 2019) and we tested for differential gene expression between strains, 3D7 versus 3D1 and 1A5 versus 1E4, using R::EdgeR (Robinson *et al*. 2010).

## Results

### Effect of KCl on growth and melanisation

A total of 158,480 and 120,059 colonies from the 1A5×1E4 and 3D7×3D1 crosses, respectively, were phenotyped (see Table S2 for details). Differences in growth and melanisation between salt stress and control environments, and growth rate KCl tolerance and melanisation rate KCl tolerance can be seen in Figure 1. We used paired Wilcoxon tests to compare mean trait values between the control and KCl treatments (Table S2). In the presence of KCl, colony radius was smaller at 8 and 12 dpi in both crosses, and the growth rate was reduced in the 3D7×3D1 cross, but not in the 1A5×1E4 cross, suggesting that isolates from the 3D7×3D1 cross were more sensitive to KCl (Figure 1, Table S2). This is consistent with the greater growth rate tolerance in the 1A5×1E4 cross compared to the 3D7×3D1 cross (*F*_(1,487)_ = 9.19, *p*=0.002, growth rate KCl tolerance: 1A5×1E4 = 1.04, 3D7×3D1 = 0.96). In the KCl environment we observed less melanisation at both time points and a similar reduced accumulation of melanin over time (melanisation rate) in both crosses (Figure 1, Table S2, *F*_(1,486)_ = 2.85, *p*=0.09, melanisation rate KCl tolerance: 1A5×1E4 = 0.196, 3D7×3D1 = 0.149).

**Figure 1.**
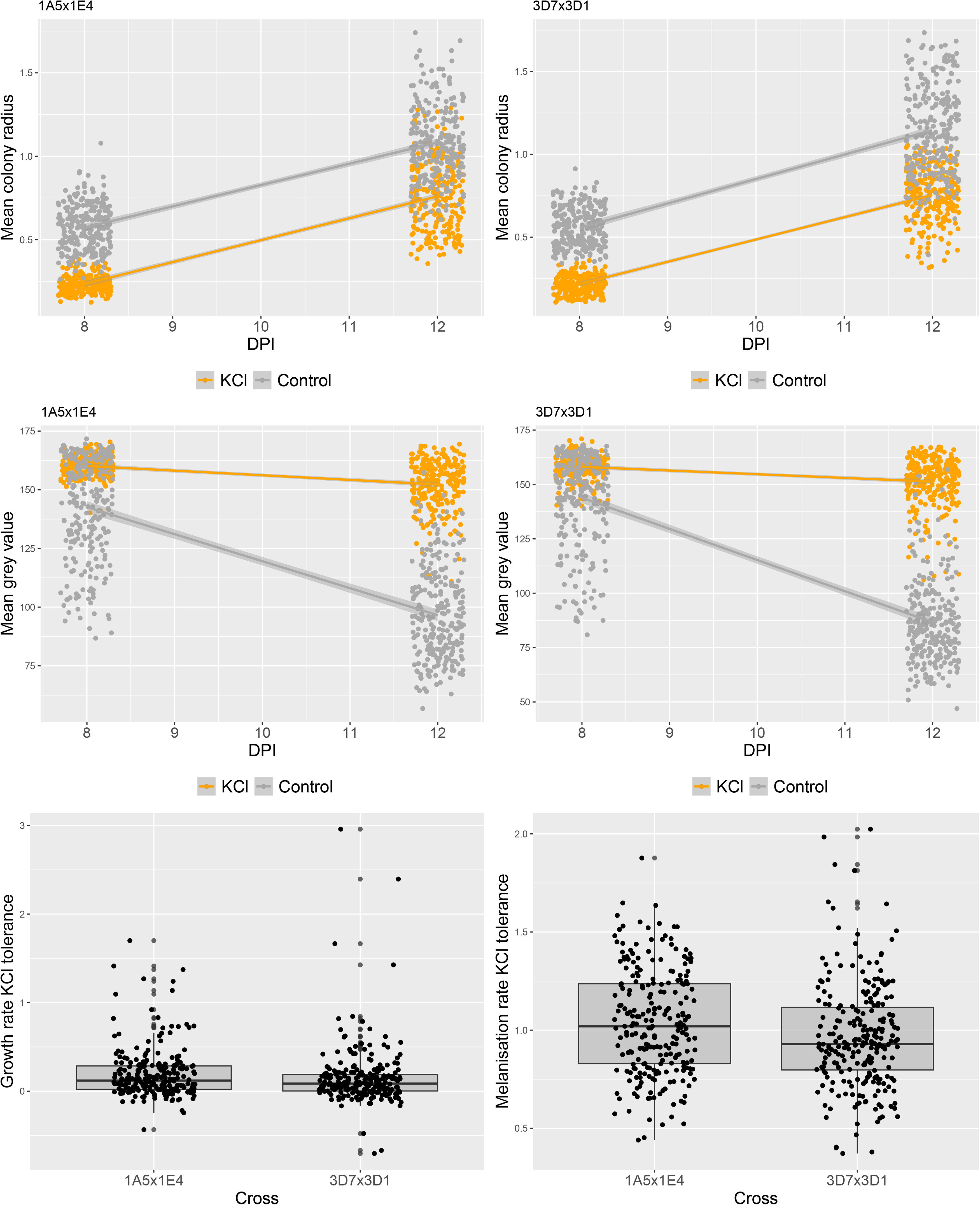
Trait variation in the control and salt stress environments across offspring in each cross. Top panel: Points are mean colony radius in the control and KCl environments at 8 and 12 dpi, smoothed lines demonstrate linear change over time. Middle panel: Points are mean grey value in the control and KCL environments at 8 and 12 dpi, smoothed lines demonstrate linear change over time. Bottom panel: Growth rate KCl tolerance and melanisation rate KCl tolerance (KCl tolerance = rate in KCl/rate in control conditions) between the two crosses.

### Correlations between traits

All pairwise correlations (Pearson, non-parametric) between independent measurements are provided in Tables S3 and S4. Excluding traits that are statistically or functionally coupled, which includes: correlations between two time points of the same trait (e.g. colony radius at 8 dpi and 12 dpi in the control), correlations between a rate and the corresponding time point measurements (e.g. grey value at 12 dpi and melanisation rate in the control), or correlations between rates and KCl tolerance measurements, we identified many correlations between independent measurements (Table S3 and S4). Notably, growth rate in the control environment was correlated with growth rate in the KCl environment in both crosses (*r*(1A5×1E4) = 0.63 and *r*(3D7×3D1) = 0.62, Figure S1) – thus isolates that grow faster in a benign environment also grow faster in the presence of salt stress i.e., they have a higher intrinsic growth rate. Melanisation rates showed a similar trend in 1A5×1E4 – isolates that had higher melanisation rates in the benign environment also had higher melanisation rates under KCl stress (Table S4, *r* = 0.19), but this was not observed in the 3D7×3D1 cross (Table S3, *r* = 0.07, Figure S1).

We were particularly interested in the relationship between KCl tolerance and melanisation, as melanin can help protect cells from osmotic stress. In both crosses, the mean grey value in the control environment at 12 dpi was negatively correlated with growth rate KCl tolerance (Tables S2 and S4, 1A5×1E4 *r* = -0.26 and 3D7×3D1 *r* = -0.43, Figure S2). The relationship at 8 dpi was also negative in both crosses, but not significant in the 3D7×3D1 cross (Tables S3 and S4, 1A5×1E4 *r* = -0.44 and 3D7×31 *r* = -0.12, Figure S2). As lower mean grey values correspond to higher amounts of melanin, these correlations indicate that isolates that produce more melanin in a benign environment have a growth rate that is more tolerant to KCl stress, although this relationship was not significant at 8 dpi in the 3D7×3D1 cross.

### QTL results

We identified at least one significant QTL for all traits in the 3D7×2D1 cross and all but four traits (melanisation rate and mean grey value at 12 dpi in the KCl environment, and melanisation rate KCl tolerance at 12dpi, Melanisation rate KCl tolerance) in the 1A5×1E4 cross (Table 1). In general, the traits lacking significant QTL had relatively low heritability (Table 1) and thus the power to detect a QTL for those traits was relatively low. All QTL positions, interval sizes and numbers of genes within the interval are presented in Tables 2 and 3. Largely overlapping QTL were grouped and given a unique QTL identifier. Examples of how traits measured in different environments can have overlapping QTL intervals are shown in Figure 2. The QTL No. 4, 9 and 11 are each regions of the genome that explain variation in growth rate in both the KCl and control environments. As expected, traits that were highly correlated and/or statistically coupled often had a QTL peak at the same, or overlapping positions (e.g. the 1A5×1E4 grey values in the control environment at 8 dpi and 12 dpi mapped to the same position on Chr 2: 1438934bp; the 3D7×3D1 QTL for colony radius at 12 dpi and growth rate in the control environment mapped to Chr 8: 1100862bp).

**Figure 2.**
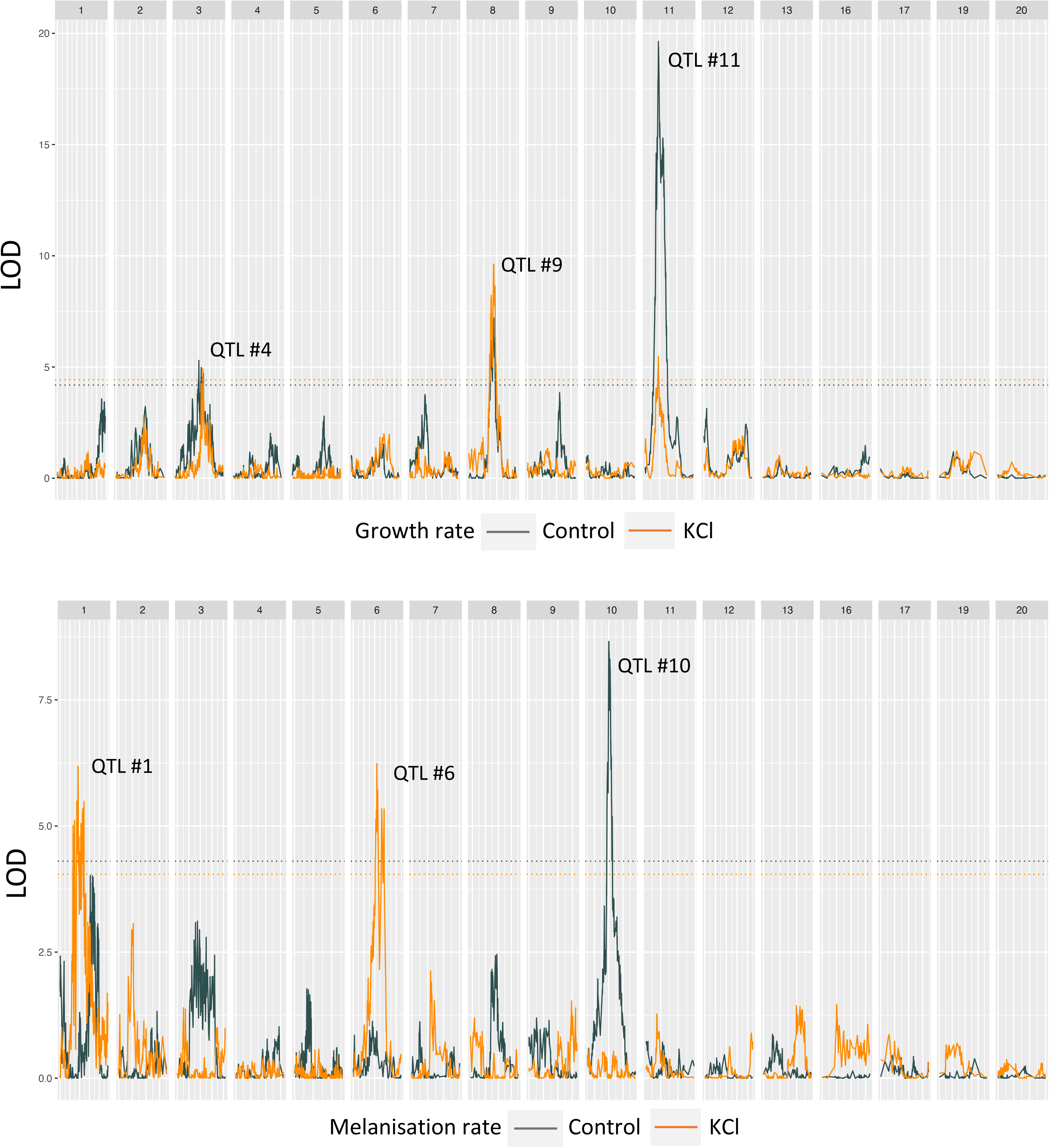
LOD plots from interval mapping for representative traits a) growth rate and b) melanisation rate in the control environment (grey) and in the KCl environment (orange) in the 3D7×3D1 cross. Each sub-panel is a chromosome with the chromosome number indicated at the top. Some numbers are missing because *Zymoseptoria tritici* carries eight accessory chromosomes (Chr 14-21) that show presence/absence polymorphisms among isolates. The dashed horizontal lines indicate the LOD significance thresholds for each trait based on permutation. The numbers above the QTL peaks correspond to the QTL numbers from Table 2. For LOD plots of all traits see Supplementary Information Figure S3.

**Table 2.**
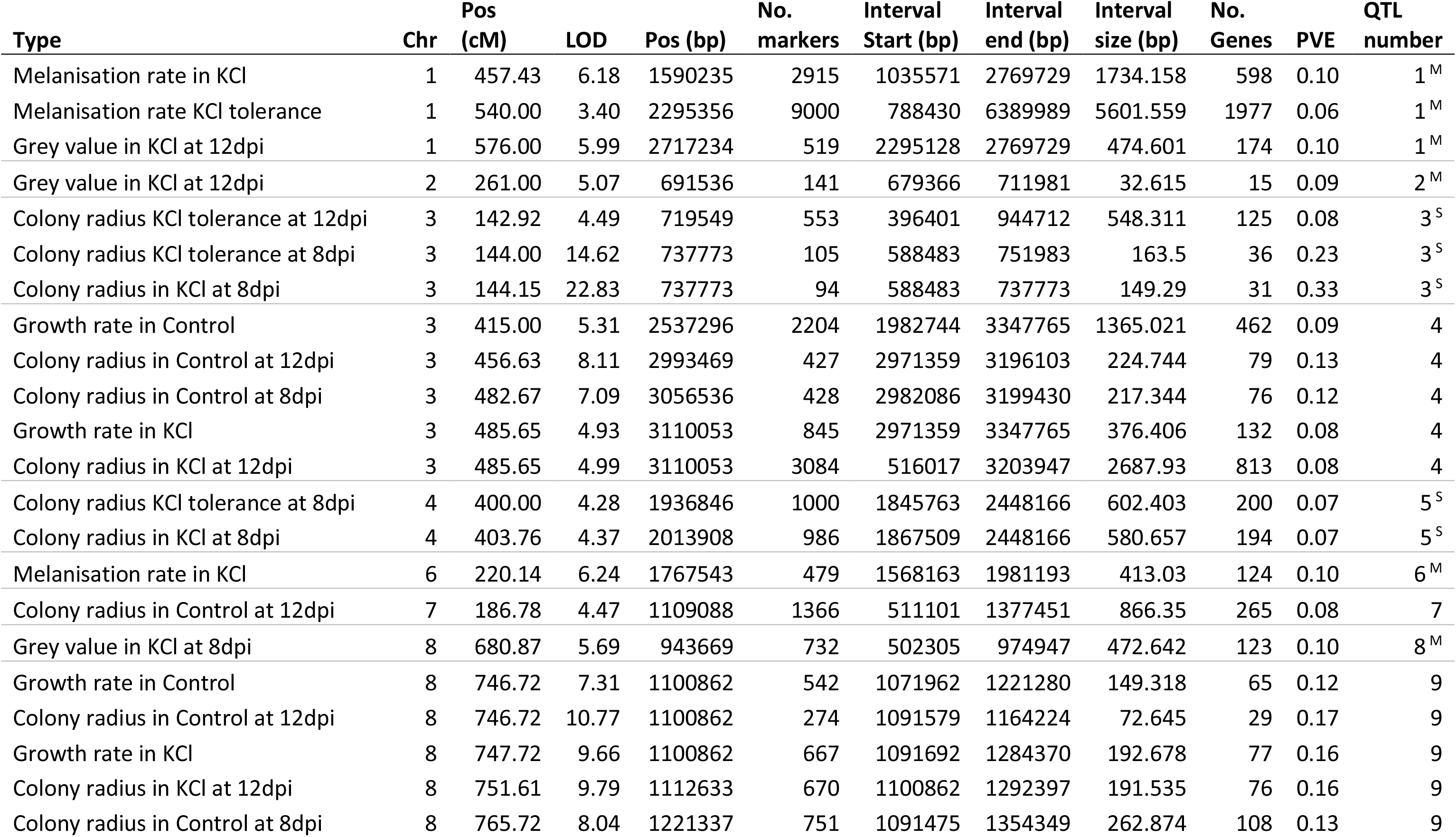
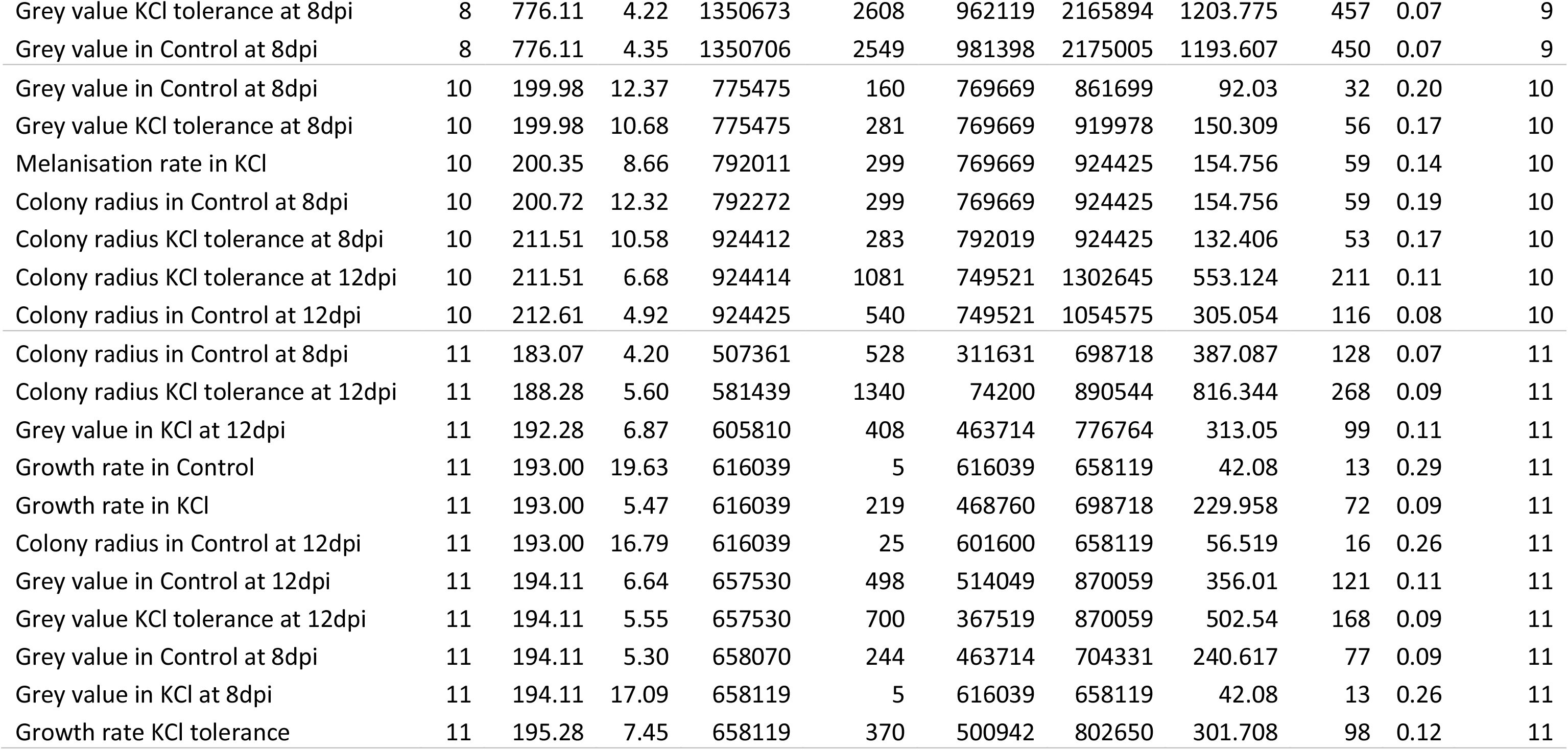
Description of all QTL identified from the 3D7×3D1 cross, including map and physical position of the LOD peak marker, the Bayes credible interval length and the number of genes within the interval. PVE is the proportion of variance explained by the QTL. QTL number is a number assigned to a particular QTL peak which groups overlapping QTLs. Superscript letters indicate QTL unique to KCL environment for size-related (^S^) and melanin-related traits (^M^).

| Type | Chr | Pos (cM) | LOD | Pos (bp) | No. markers | Interval Start (bp) | Interval end (bp) | Interval size (bp) | No. Genes | PVE | QTL number |
| --- | --- | --- | --- | --- | --- | --- | --- | --- | --- | --- | --- |
| Melanisation rate in KCL | 1 | 457.43 | 6.18 | 1590235 | 2915 | 1035571 | 2769729 | 1734.158 | 598 | 0.10 | 1<sup>M</sup> |
| Melanisation rate KCL tolerance | 1 | 540.00 | 3.40 | 2295356 | 9000 | 788430 | 6389989 | 5601.559 | 1977 | 0.06 | 1<sup>M</sup> |
| Grey value in KCL at 12dpi | 1 | 576.00 | 5.99 | 2717234 | 519 | 2295128 | 2769729 | 474.601 | 174 | 0.10 | 1<sup>M</sup> |
| Grey value in KCL at 12dpi | 2 | 261.00 | 5.07 | 691536 | 141 | 679366 | 711981 | 32.615 | 15 | 0.09 | 2<sup>M</sup> |
| Colony radius KCL tolerance at 12dpi | 3 | 142.92 | 4.49 | 719549 | 553 | 396401 | 944712 | 548.311 | 125 | 0.08 | 3<sup>S</sup> |
| Colony radius KCL tolerance at 8dpi | 3 | 144.00 | 14.62 | 737773 | 105 | 588483 | 751983 | 163.5 | 36 | 0.23 | 3<sup>S</sup> |
| Colony radius in KCL at 8dpi | 3 | 144.15 | 22.83 | 737773 | 94 | 588483 | 737773 | 149.29 | 31 | 0.33 | 3<sup>S</sup> |
| Growth rate in Control | 3 | 415.00 | 5.31 | 2537296 | 2204 | 1982744 | 3347765 | 1365.021 | 462 | 0.09 | 4 |
| Colony radius in Control at 12dpi | 3 | 456.63 | 8.11 | 2993469 | 427 | 2971359 | 3196103 | 224.744 | 79 | 0.13 | 4 |
| Colony radius in Control at 8dpi | 3 | 482.67 | 7.09 | 3056536 | 428 | 2982086 | 3199430 | 217.344 | 76 | 0.12 | 4 |
| Growth rate in KCL | 3 | 485.65 | 4.93 | 3110053 | 845 | 2971359 | 3347765 | 376.406 | 132 | 0.08 | 4 |
| Colony radius in KCL at 12dpi | 3 | 485.65 | 4.99 | 3110053 | 3084 | 516017 | 3203947 | 2687.93 | 813 | 0.08 | 4 |
| Colony radius KCL tolerance at 8dpi | 4 | 400.00 | 4.28 | 1936846 | 1000 | 1845763 | 2448166 | 602.403 | 200 | 0.07 | 5<sup>S</sup> |
| Colony radius in KCL at 8dpi | 4 | 403.76 | 4.37 | 2013908 | 986 | 1867509 | 2448166 | 580.657 | 194 | 0.07 | 5<sup>S</sup> |
| Melanisation rate in KCL | 6 | 220.14 | 6.24 | 1767543 | 479 | 1568163 | 1981193 | 413.03 | 124 | 0.10 | 6<sup>M</sup> |
| Colony radius in Control at 12dpi | 7 | 186.78 | 4.47 | 1109088 | 1366 | 511101 | 1377451 | 866.35 | 265 | 0.08 | 7 |
| Grey value in KCL at 8dpi | 8 | 680.87 | 5.69 | 943669 | 732 | 502305 | 974947 | 472.642 | 123 | 0.10 | 8<sup>M</sup> |
| Growth rate in Control | 8 | 746.72 | 7.31 | 1100862 | 542 | 1071962 | 1221280 | 149.318 | 65 | 0.12 | 9 |
| Colony radius in Control at 12dpi | 8 | 746.72 | 10.77 | 1100862 | 274 | 1091579 | 1164224 | 72.645 | 29 | 0.17 | 9 |
| Growth rate in KCL | 8 | 747.72 | 9.66 | 1100862 | 667 | 1091692 | 1284370 | 192.678 | 77 | 0.16 | 9 |
| Colony radius in KCL at 12dpi | 8 | 751.61 | 9.79 | 1112633 | 670 | 1100862 | 1292397 | 191.535 | 76 | 0.16 | 9 |
| Colony radius in Control at 8dpi | 8 | 765.72 | 8.04 | 1221337 | 751 | 1091475 | 1354349 | 262.874 | 108 | 0.13 | 9 |
| Grey value KCl tolerance at 8dpi | 8 | 776.11 | 4.22 | 1350673 | 2608 | 962119 | 2165894 | 1203.775 | 457 | 0.07 | 9 |
| Grey value in Control at 8dpi | 8 | 776.11 | 4.35 | 1350706 | 2549 | 981398 | 2175005 | 1193.607 | 450 | 0.07 | 9 |
| Grey value in Control at 8dpi | 10 | 199.98 | 12.37 | 775475 | 160 | 769669 | 861699 | 92.03 | 32 | 0.20 | 10 |
| Grey value KCl tolerance at 8dpi | 10 | 199.98 | 10.68 | 775475 | 281 | 769669 | 919978 | 150.309 | 56 | 0.17 | 10 |
| Melanisation rate in KCl | 10 | 200.35 | 8.66 | 792011 | 299 | 769669 | 924425 | 154.756 | 59 | 0.14 | 10 |
| Colony radius in Control at 8dpi | 10 | 200.72 | 12.32 | 792272 | 299 | 769669 | 924425 | 154.756 | 59 | 0.19 | 10 |
| Colony radius KCl tolerance at 8dpi | 10 | 211.51 | 10.58 | 924412 | 283 | 792019 | 924425 | 132.406 | 53 | 0.17 | 10 |
| Colony radius KCl tolerance at 12dpi | 10 | 211.51 | 6.68 | 924414 | 1081 | 749521 | 1302645 | 553.124 | 211 | 0.11 | 10 |
| Colony radius in Control at 12dpi | 10 | 212.61 | 4.92 | 924425 | 540 | 749521 | 1054575 | 305.054 | 116 | 0.08 | 10 |
| Colony radius in Control at 8dpi | 11 | 183.07 | 4.20 | 507361 | 528 | 311631 | 698718 | 387.087 | 128 | 0.07 | 11 |
| Colony radius KCl tolerance at 12dpi | 11 | 188.28 | 5.60 | 581439 | 1340 | 74200 | 890544 | 816.344 | 268 | 0.09 | 11 |
| Grey value in KCl at 12dpi | 11 | 192.28 | 6.87 | 605810 | 408 | 463714 | 776764 | 313.05 | 99 | 0.11 | 11 |
| Growth rate in Control | 11 | 193.00 | 19.63 | 616039 | 5 | 616039 | 658119 | 42.08 | 13 | 0.29 | 11 |
| Growth rate in KCl | 11 | 193.00 | 5.47 | 616039 | 219 | 468760 | 698718 | 229.958 | 72 | 0.09 | 11 |
| Colony radius in Control at 12dpi | 11 | 193.00 | 16.79 | 616039 | 25 | 601600 | 658119 | 56.519 | 16 | 0.26 | 11 |
| Grey value in Control at 12dpi | 11 | 194.11 | 6.64 | 657530 | 498 | 514049 | 870059 | 356.01 | 121 | 0.11 | 11 |
| Grey value KCl tolerance at 12dpi | 11 | 194.11 | 5.55 | 657530 | 700 | 367519 | 870059 | 502.54 | 168 | 0.09 | 11 |
| Grey value in Control at 8dpi | 11 | 194.11 | 5.30 | 658070 | 244 | 463714 | 704331 | 240.617 | 77 | 0.09 | 11 |
| Grey value in KCl at 8dpi | 11 | 194.11 | 17.09 | 658119 | 5 | 616039 | 658119 | 42.08 | 13 | 0.26 | 11 |
| Growth rate KCl tolerance | 11 | 195.28 | 7.45 | 658119 | 370 | 500942 | 802650 | 301.708 | 98 | 0.12 | 11 |

**Table 3.** Description of all QTLs identified from the 1A5×1E4 cross, including map and physical position of the LOD peak marker, the Bayes credible interval length and the number of genes within the interval. PVE is the proportion of variance explained by the QTL. QTL number is a number assigned to a particular QTL peak which identifies overlapping QTLs. Superscript letters indicate QTL unique to KCL environment for size-related (^S^) and melanin-related traits (^M^).

| Type | Chr | Pos (cM) | LOD | Pos (bp) | No. markers | Interval Start (bp) | Interval end (bp) | Interval size (bp) | No. Genes | PVE | QTL number |
| --- | --- | --- | --- | --- | --- | --- | --- | --- | --- | --- | --- |
| Grey value in Control at 8dpi | 1 | 328.91 | 5.69 | 572065 | 402 | 442697 | 727696 | 284.999 | 92 | 0.10 | 1 |
| Grey value KCl tolerance at 8dpi | 1 | 328.91 | 5.01 | 572065 | 405 | 442697 | 732023 | 289.326 | 93 | 0.08 | 1 |
| Melanisation rate KCl tolerance | 1 | 328.91 | 4.46 | 572065 | 463 | 461577 | 779936 | 318.359 | 102 | 0.08 | 1 |
| Colony radius KCl tolerance at 12dpi | 1 | 559.12 | 4.07 | 1849200 | 4654 | 399579 | 5725682 | 5326.103 | 1964 | 0.07 | 2<sup>S</sup> |
| Growth rate KCl tolerance | 1 | 797.00 | 4.80 | 3405315 | 3068 | 399544 | 3488556 | 3089.012 | 1151 | 0.08 | 2<sup>S</sup> |
| Colony radius in KCl at 12dpi | 1 | 1104.00 | 4.03 | 5792644 | 4774 | 372886 | 5956797 | 5583.911 | 2056 | 0.07 | 2<sup>S</sup> |
| Melanisation rate in Control | 2 | 70.00 | 6.56 | 574965 | 1211 | 405115 | 1914465 | 1509.35 | 509 | 0.11 | 3 |
| Grey value in Control at 12dpi | 2 | 204.66 | 4.55 | 1438934 | 618 | 1284536 | 2152463 | 867.927 | 281 | 0.08 | 4 |
| Grey value in Control at 8dpi | 2 | 204.66 | 18.08 | 1438934 | 185 | 1438863 | 1748215 | 309.352 | 95 | 0.27 | 4 |
| Grey value KCl tolerance at 8dpi | 2 | 204.66 | 15.97 | 1438934 | 184 | 1438934 | 1748215 | 309.281 | 95 | 0.24 | 4 |
| Colony radius in Control at 8dpi | 2 | 211.42 | 8.76 | 1631840 | 217 | 1438934 | 1776132 | 337.198 | 105 | 0.14 | 4 |
| Grey value KCl tolerance at 8dpi | 3 | 147.00 | 6.15 | 1325440 | 210 | 1243932 | 1684531 | 440.599 | 179 | 0.10 | 5 |
| Grey value in Control at 8dpi | 3 | 148.00 | 6.66 | 1325440 | 133 | 1325422 | 1592801 | 267.379 | 115 | 0.11 | 5 |
| Colony radius in Control at 8dpi | 3 | 163.74 | 5.83 | 1521535 | 725 | 355042 | 1592801 | 1237.759 | 374 | 0.10 | 5 |
| Melanisation rate in Control | 3 | 193.38 | 11.99 | 1711037 | 290 | 1489966 | 2105601 | 615.635 | 180 | 0.19 | 6 |
| Grey value in Control at 12dpi | 3 | 198.73 | 4.04 | 1928783 | 860 | 1743111 | 2927356 | 1184.245 | 361 | 0.07 | 6 |
| Growth rate KCl tolerance | 5 | 399.00 | 4.34 | 2638913 | 155 | 2511016 | 2658480 | 147.464 | 48 | 0.07 | 7<sup>S</sup> |
| Colony radius in KCl at 8dpi | 6 | 179.00 | 4.48 | 1512508 | 594 | 778583 | 1557528 | 778.945 | 256 | 0.08 | 8<sup>S</sup> |
| Growth rate in Control | 7 | 158.82 | 4.43 | 539824 | 255 | 437699 | 651318 | 213.619 | 80 | 0.07 | 9 |
| Colony radius in Control at 12dpi | 7 | 158.82 | 4.67 | 539837 | 254 | 437718 | 651318 | 213.6 | 80 | 0.08 | 9 |
| Growth rate in KCl | 8 | 76.80 | 8.79 | 274792 | 82 | 256529 | 346224 | 89.695 | 25 | 0.14 | 10 |
| Colony radius in KCl at 12dpi | 8 | 76.80 | 8.94 | 274792 | 67 | 256529 | 313758 | 57.229 | 17 | 0.15 | 10 |
| Grey value in Control at 12dpi | 8 | 78.58 | 4.92 | 302772 | 882 | 89684 | 921404 | 831.72 | 308 | 0.08 | 10 |
| Grey value in Control at 8dpi | 8 | 79.00 | 12.12 | 302772 | 66 | 270834 | 346224 | 75.39 | 22 | 0.19 | 10 |
| Colony radius in Control at 12dpi | 8 | 79.00 | 9.72 | 302772 | 80 | 260355 | 346224 | 85.869 | 24 | 0.16 | 10 |
| Colony radius in Control at 8dpi | 8 | 79.00 | 11.06 | 302772 | 415 | 260101 | 732444 | 472.343 | 163 | 0.18 | 10 |
| Grey value KCl tolerance at 8dpi | 8 | 79.00 | 8.26 | 302772 | 339 | 89573 | 346224 | 256.651 | 90 | 0.14 | 10 |
| Colony radius KCl tolerance at 8dpi | 8 | 137.00 | 5.05 | 832180 | 1177 | 589084 | 1648406 | 1059.322 | 396 | 0.08 | 10 |
| Grey value in KCl at 8dpi | 8 | 164.00 | 4.33 | 1377926 | 1406 | 305075 | 1623437 | 1318.362 | 485 | 0.07 | 10 |
| Melanisation rate in Control | 12 | 126.74 | 4.24 | 473688 | 275 | 441706 | 821603 | 379.897 | 101 | 0.07 | 11 |

LOD plots for all traits listed in Tables 2 and 3 are provided in the Supplementary Figures S4 and Figure S5. In Figures 2 and 3 we provide example plots for only four traits (growth rate in KCl, growth rate in the control, melanisation rate in KCl and melanisation rate in the control) for the two crosses.

**Figure 3.**
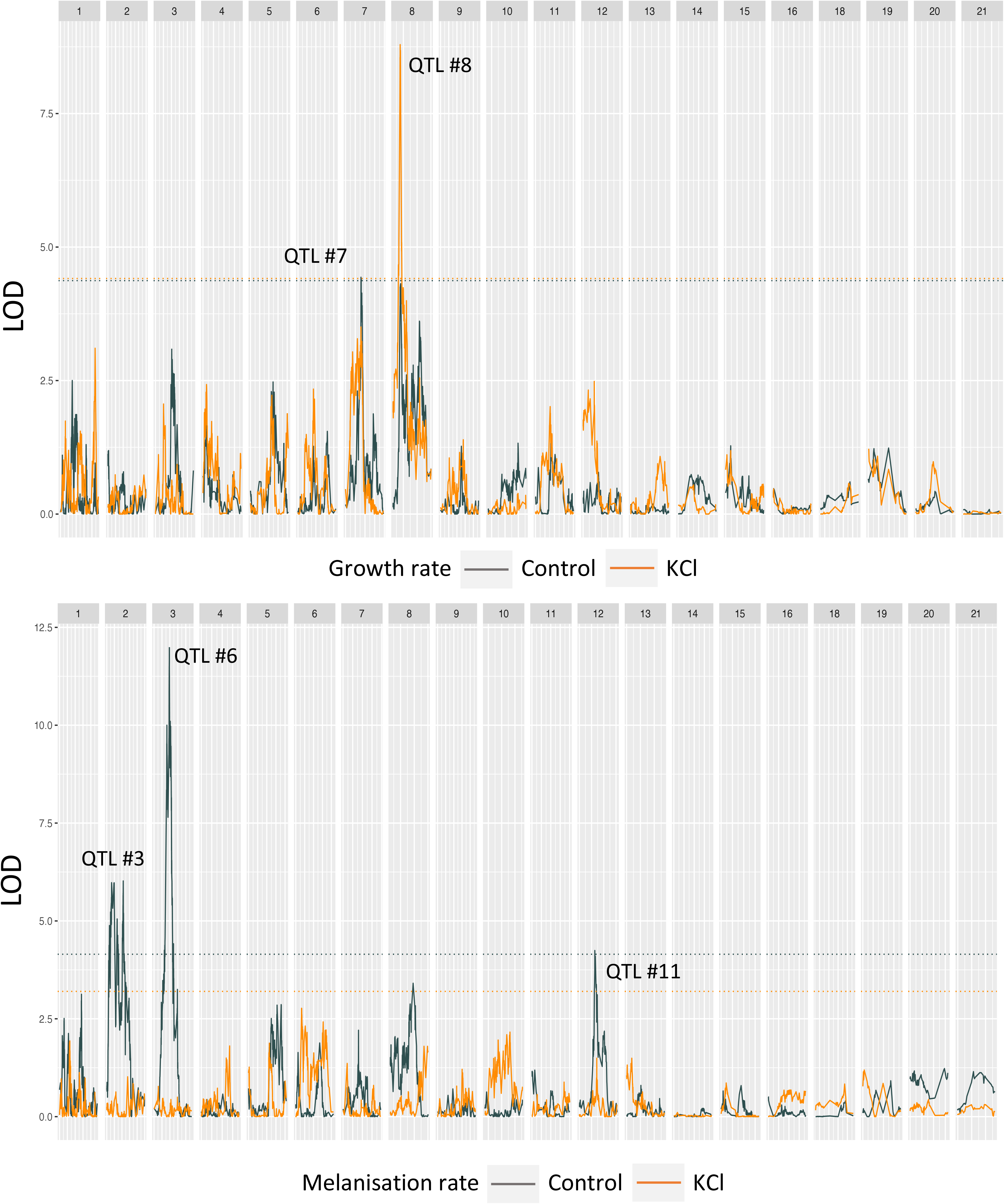
LOD plots from interval mapping for representative traits a) growth rate and b) melanisation rate in the control environment (grey) and in the KCl environment (orange) in the 1A5×1E4 cross. Some numbers are missing because *Zymoseptoria tritici* carries eight accessory chromosomes (Chr 14-21) that show presence/absence polymorphisms among isolates. The dashed horizontal lines indicate the LOD significance thresholds for each trait based on permutation. The numbers above the QTL peaks correspond to the QTL numbers from Table 3. For LOD plots of all traits see Supplementary Figure S4.

### QTL shared across environments, traits, and time points

Several QTL were shared amongst benign and salt-stress environments, across time points and traits. Only one of these shared QTL is novel to this study, the others were found in earlier studies investigating different stressors. The single novel QTL shared between benign and salt-stress environments was found in the 3D7×3D1 cross on Chr 8 (1100.86-1221.33 Kb, QTL No. 9, Figure 2) and explained variation in seven traits including growth in the control and KCl environments, as well as mean grey value in the control environment and grey value KCl tolerance at 8 dpi (Figure 2, Table 2).

Of the QTL shared across benign and KCl environments that were identified here and found in earlier studies using different stresses (e.g. cold, fungicide), the most striking – in terms of the number of traits and the magnitude of the LODs, was the QTL on Chr 11 between 507.36-658.11 Kb (QTL No. 12) in the 3D7×3D1 cross. This QTL explained variation in 11 of the 18 traits, and in both environments (Table 2). This QTL was identified previously and was found to explain variation in growth, fungicide sensitivity and melanisation in this cross (Lendenmann *et al*. 2014, 2015). The next largest, in terms of number of traits, was a QTL in the 1A5×1E4 cross on Chr 8 between 89.573-346.224 Kb (QTL No. 10) that explained variation in growth rate, colony radius and grey value in both control and KCl environments (Figure 3, Table 3). This QTL on Chr 8 was also found to explain variation in response to oxidative stress (Zhong *et al*. 2021). Another notable QTL in terms of high LOD (>10) was on Chr 10 (749.521– 1302.645 Kb, QTL No. 10) in the 3D7×3D1 cross, which explained variation in melanisation rate and colony radius in the control environment (Table 2). This QTL was previously found to explain variation in growth-related traits at 10^0^C (Lendenmann *et al*. 2014) and under oxidative stress (Zhong *et al*. 2021).

### Unique QTL associated with the KCl environment and pathway enrichment analysis

Several novel QTL that were unique to the KCl environment were also found (indicated with an ^S or M^ in Table 2, 3). In the 3D7×3D1 cross, two QTL were found for traits related to colony radius: the first on Chr 3 (QTL No. 3) explained variation in colony radius in KCl at 8 dpi and colony radius KCl tolerance at 8 and 12 dpi; and the second on Chr 4 (QTL No. 5) explained variation in colony radius in KCl at 8 dpi and colony radius KCl tolerance at 8 dpi. For melanin-related traits, four QTL were found in the 3D7×3D1 cross: the first on Chr 1 (QTL No. 1) explained variation in three traits (melanisation rate in KCl, melanisation rate KCl tolerance and grey value in KCL at 12 dpi); the second on Chr 2 (QTL No. 2) explained variation in grey value in KCl at 12 dpi; the third on Chr 6 (QTL No. 6) explained variation in melanisation rate in KCl; and the fourth on Chr 8 (QTL No. 8) explained variation in grey value in KCl at 8 dpi. In the 1A5×1E4 cross a total of three QTL related to colony radius in KCl were found. The first was on Chr 1: QTL No. 2 explained variation in colony radius KCl tolerance at 12 dpi, growth rate KCl tolerance, and colony radius in KCl at 12 dpi. The second unique KCl environment QTL was on Chr 5 (QTL No. 7) and explained variation in growth rate KCl tolerance; the fifth on Chr 6 (QTL No. 8) explained variation in radius at 8 dpi in KCl. No unique QTL were found for melanin-related traits in the 1A5×1E4 cross.

For pathway enrichment analyses we focussed on traits that had significant QTLs unique to the KCl environment (indicated in Tables 2 and 3 with ^S or M^) and investigated if genes associated with those traits were enriched for KEGG pathways or GO annotations. We analysed each trait-QTL separately and did the same analysis using all genes for size- or melanin-related traits combined (Table S5, Table S6). No significantly enriched KEGG pathways were identified in our data. For the GO enrichment we report on the results for size- or melanin-related trait-QTL combined and we use the raw p-values instead of the adjusted p-values as indicators of possible GO enrichment because adjusting p-values in GO enrichment analysis may be too conservative and inappropriate (Alexa and Rahnenfuhrer 2023). All results are presented in Supplementary Tables S5 and S6. Considering all genes within size-related unique KCl QTL combined in the 3D7×3D1 cross (Table 2, QTL No.^S^): for Biological Process (BP) double-strand break repair via nonhomologous end joining (GO:0006303) and nucleotide-excision repair (GO:0006289) GO terms were enriched; for Cellular Component (CC) Ku70:Ku80 complex (GO:0043564) GO term was enriched; and for Molecular Function (MF) GTPase activity (GO:0003924), superoxide dismutase activity (GO:0004784), telomeric DNA binding (GO:0042162) and damaged DNA binding (GO:0003684) GO terms were enriched (Table S5b). There were 15 genes within these QTLs that have one or more of these GO annotations (Table S7a). For genes within melanin-related unique KCl QTL combined in the 3D7×3D1 cross (Table 2 QTL No.^M^): for BP glutathione biosynthetic process (GO:0006750) GO terms was enriched; for CC no GO term was enriched; and for MF chaperone binding (GO:0051087), phosphatidylinositol phospholipase C activity (GO:0004435), and shikimate 3-dehydrogenase (NADP+) activity (GO:0004764) GO terms were enriched (Table S5d). There were 13 genes within these QTLs that have one or more of these GO annotations (Table S7b). Considering all genes within size-related unique KCl QTL combined in the 1A5×1E4 cross (Table 2 QTL No.^S^): for BP no GO terms were enriched; for CC cell wall (GO:0005618) GO term was enriched; and for MF chaperone binding (GO:0051087) and triglyceride lipase activity (GO:0004806) GO terms were enriched (Table S6b). There were 12 genes within these QTLs that have one or more of these GO annotations (Table S7c). There were no unique KCl melanin-related QTL in the 1A5×1E4 cross.

### Analysis of QTL No. 3 in 3D7×3D1 and identification of candidate genes

To identify candidate genes associated with KCl stress, we focused attention on novel QTL with a high LOD (>10), relatively narrow intervals, and that explained a relatively high proportion of the variation for a KCl-associated trait. Only QTL No. 3 in the 3D7×3D1 cross met these requirements (Table 2). This QTL had a high LOD (>10) for colony radius at 8 dpi in the KCl environment and colony radius KCl tolerance at 8 dpi. The LOD peaked at 22.8 at chromosome position 737773 bp. This QTL explained 33% of the phenotypic variance in colony radius at 8 dpi and 23% of the variance in colony radius KCl tolerance at 8 dpi. Isolates with the non-reference allele at the QTL peak marker had larger colonies at 8 dpi (*F*=113.1, df=1,233, *p*=<0.001, Figure 4c) and greater colony radius KCl tolerance at 8 dpi (*F*=56.07, df=1,23, *p*=<0.001). The Bayes confidence interval for the QTL for colony radius in KCl at 8 dpi spanned 149.29 Kb and contained 94 SNPs and 36 genes (Table 2, Figure 4b). For this interval we present data on the genes in the reference genome (3D7), the orthologous genes in 3D1, the gene sequence similarity (including ±200bp of flanking regions) between the parents, the gene’s putative encoded function (PFAM database) and if there is any evidence of differential gene expression between the parents across all available datasets (Table 4 and see Table S8 for full details).

**Figure 4.**
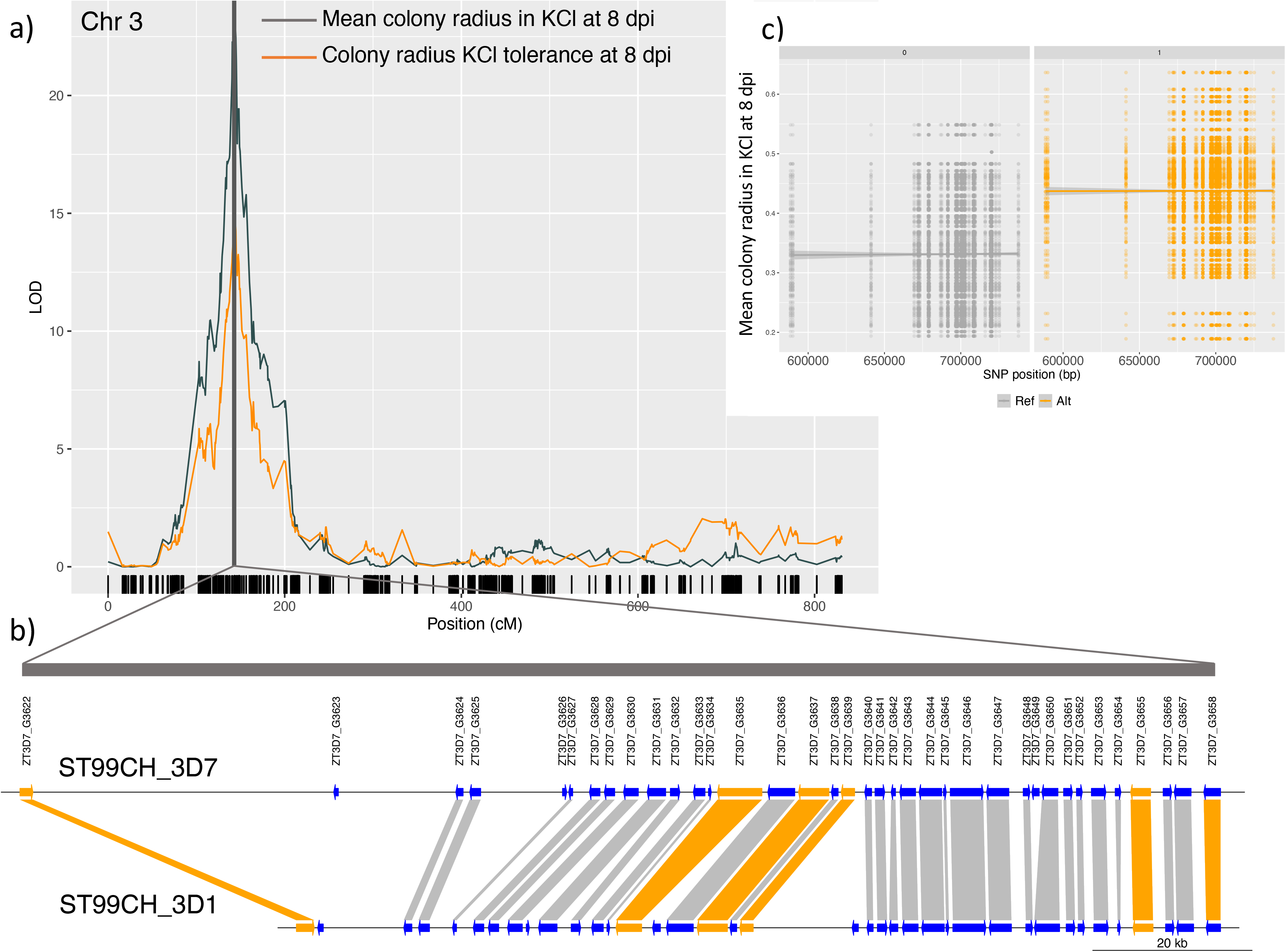
QTL on Chromosome 3 for three traits related to growth under osmotic stress in the 3D7×3D1 cross. The three traits were colony radius at 8 dpi, colony radius KCl tolerance at 8 and 12 dpi (KCl tolerance = colony radius in KCl/colony radius in control conditions). A) LOD peak and Bayes credible interval (shaded) from interval mapping, with SNP positions indicated on the x-axis as black vertical lines, b) Alignment of the 36 genes within the QTL interval between the 3D7 and 3D1 parental strains – orange-coloured genes are the six most promising candidate genes described in the main text and c) allelic effects of 94 SNPs within the QTL interval on mean colony radius in KCl at 8 dpi. Isolates carrying the reference allele (ref=3D7) shown in grey have smaller colonies compared to the alternative allele (alt=3D1) shown in orange. SNP position in base pairs is shown on the x-axis. The marker with the highest LOD is at position 737773 bp.

**Table 4.** Genes within the QTL interval for colony radius at 8 dpi in KCl and colony radius KCl tolerance at 8 dpi in the 3D7×3D1 cross. Bold indicates genes that are the most promising candidate genes, as described in the main text. Gene sequence similarity between the two parental strains was obtained using BLAST. DGE: Differential gene expression column indicates if the gene was differentially expressed between parents in any one of the data sets used (in planta or in vitro, see Methods in main text). The annotations were obtained using Interproscan using multiple protein databases, here we show only the PFAM annotation.

| Chr | Gene id 3D7 | Gene start | Gene end | Gene id 3D1 | Sequence similarity (%) | DGE | Annotation |
| --- | --- | --- | --- | --- | --- | --- | --- |
| **3** | **ZT3D7_G3622** | **598091** | **599743** | **ZT3D1_G3567** | **97.68** | **yes** | **Fungal-specific transcription factor domain** |
| 3 | ZT3D7_G3623 | 637954 | 638542 | ZT3D1_G7387 | 96.23 | NA |  |
| 3 | ZT3D7_G3624 | 653468 | 654397 | ZT3D1_G3569 | 90.78 | yes | Protein of unknown function (DUF1349) |
| 3 | ZT3D7_G3625 | 655237 | 656628 | ZT3D1_G3570 | 97.04 | yes |  |
| 3 | ZT3D7_G3626 | 667030 | 667567 | NA |  | yes | Pterin 4 alpha carbinolamine dehydratase |
| 3 | ZT3D7_G3627 | 667812 | 668321 | ZT3D1_G3571 | 98.46 | yes |  |
| 3 | ZT3D7_G3628 | 670445 | 671787 | ZT3D1_G3572 | 100.00 | yes | Transketolase, pyrimidine binding domain, Transketolase, C-terminal domain |
| 3 | ZT3D7_G3629 | 672336 | 673658 | ZT3D1_G3573 | 99.83 | NA |  |
| 3 | ZT3D7_G3630 | 674777 | 676658 | ZT3D1_G3574 | 99.96 | yes | Dehydratase family |
| 3 | ZT3D7_G3631 | 677782 | 680135 | ZT3D1_G3576 | 97.72 | yes | ABC transporter, ABC transporter |
| 3 | ZT3D7_G3632 | 680734 | 681942 | ZT3D1_G3577 | 97.73 | yes | Epoxide hydrolase N terminus |
| 3 | ZT3D7_G3633 | 683647 | 685131 | ZT3D1_G3578 | 81.12 | NA | Phosphotransferase enzyme family |
| 3 | ZT3D7_G3634 | 685575 | 685937 | ZT3D1_G3579 | 98.96 | yes |  |
| **3** | **ZT3D7_G3635** | **686721** | **692363** | **ZT3D1_G3580** | **99.07** | **yes** | **Protein kinase domain, Protein kinase C terminal domain, N-terminal half of MaoC dehydratase, MaoC like domain, C2 domain** |
| 3 | ZT3D7_G3636 | 693114 | 696535 | ZT3D1_G3582 | 99.50 | yes | Importin-beta N-terminal domain, Cse1 |
| **3** | **ZT3D7_G3637** | **697014** | **700871** | **ZT3D1_G3583** | **97.44** | **yes** | **DEAD/DEAH box helicase, rRNA-processing arch domain, DSHCT (NUC185) domain, Ski2 N-terminal region, Helicase conserved C-terminal domain** |
| 3 | ZT3D7_G3638 | 701182 | 702047 | ZT3D1_G3584 | 98.74 | yes | short chain dehydrogenase |
| **3** | **ZT3D7_G3639** | **702431** | **704143** | **ZT3D1_G3585** | **97.16** | **yes** | **Major Facilitator Superfamily** |
| 3 | ZT3D7_G3640 | 705425 | 706303 | ZT3D1_G3587 | 99.91 | yes |  |
| 3 | ZT3D7_G3641 | 706740 | 707975 | ZT3D1_G3588 | 98.53 | yes |  |
| 3 | ZT3D7_G3642 | 708779 | 709336 | ZT3D1_G3589 | 98.33 | NS |  |
| 3 | ZT3D7_G3643 | 709874 | 711886 | ZT3D1_G3590 | 94.46 | yes | Ubiquitin-conjugating enzyme |
| 3 | ZT3D7_G3644 | 712327 | 715212 | ZT3D1_G3591 | 98.93 | yes | Utp14 protein |
| 3 | ZT3D7_G3645 | 715430 | 715798 | ZT3D1_G3592 | 99.74 | no | Mitochondrial ribosomal protein L27 |
| 3 | ZT3D7_G3646 | 716302 | 720497 | ZT3D1_G3593 | 98.96 | NA | Staphylococcal nuclease homologue, short chain dehydrogenase, Enoyl-<br>(Acyl carrier protein) reductase, Tudor domain |
| 3 | ZT3D7_G3647 | 720900 | 723737 | ZT3D1_G3594 | 99.38 | yes | PQQ-like domain, ER membrane protein complex subunit 1, C-terminal |
| 3 | ZT3D7_G3648 | 725571 | 726437 | ZT3D1_G3595 | 99.02 | no | Ras family |
| 3 | ZT3D7_G3649 | 726651 | 727599 | NA |  | yes | NmrA-like family |
| 3 | ZT3D7_G3650 | 727935 | 729979 | ZT3D1_G3596 | 98.53 | no | Heterokaryon incompatibility protein (HET) |
| 3 | ZT3D7_G3651 | 730698 | 731819 | ZT3D1_G3597 | 99.93 | yes | Transaldolase/Fructose-6-phosphate aldolase |
| 3 | ZT3D7_G3652 | 732329 | 733083 | ZT3D1_G3598 | 96.53 | yes | Copper/zinc superoxide dismutase (SODC) |
| 3 | ZT3D7_G3653 | 734230 | 736078 | ZT3D1_G3599 | 99.73 | yes | FGGY family of carbohydrate kinases, C-terminal domain, FGGY family of<br>carbohydrate kinases, N-terminal domain |
| 3 | ZT3D7_G3654 | 737260 | 737987 | ZT3D1_G3600 | 99.87 | yes |  |
| **3** | **ZT3D7_G3655** | **739211** | **741757** | **ZT3D1_G3601** | **99.97** | **yes** | **CBS domain, Voltage gated chloride channel** |
| 3 | ZT3D7_G3656 | 743352 | 744442 | ZT3D1_G3602 | 99.73 | yes | Protein of unknown function (DUF1295) |
| 3 | ZT3D7_G3657 | 744744 | 746896 | ZT3D1_G3603 | 99.92 | yes | Pro-kumamolisin, activation domain |
| **3** | **ZT3D7_G3658** | **748505** | **750639** | **ZT3D1_G3604** | **100** | **yes** | **PX domain, Vps5 C terminal like** |

Several genes had SNP variation predicted to have a high impact based on SNPEff analysis (Table S9, *ZT3D7_G3622*, *ZT3D7_G3623*, *ZT3D7_G3624, ZT3D7_G3632, ZT3D7_G3633, ZT3D7_G3635, ZT3D7_G3649*). A transposable element (6927 bp) was found within 200 bp of *ZT3D7_G3623*, and one gene, *ZT3D7_G3649* is a singleton – with no orthologue present in any of the other 18 *Z. tritici* reference genome sequences (Table S8). Two genes within this QTL interval and another gene immediately adjacent to the interval have GO annotations that were enriched (ZT3D7_G3648, ZT3D7_G3652, ZT3D7_G3659, Table S7). ZT3D7_G3648 and ZT3D7_G3659 have predicted GTPase activity (GO:0003924) and ZT3D7_G3652 has predicted superoxide dismutase activity (GO:0004784). The SNP variation in these genes between the parents was not predicted to have a large impact on the protein (Table S9). Based on predicted protein function, and sequence and gene expression variation between the parents, we consider six genes to be the most promising candidates for further analyses: *ZT3D7_G3622*, *ZT3D7_G3635*, *ZT3D7_G3637*, *ZT3D7_G3639*, *ZT3D7_G3655* and *ZT3D7_G3658* (this order reflects their relative positions on chromosome 3).

*ZT3D7_G3622* has a loss of function SNP (Chr 3:598092) in 3D1 that is predicted to cause a loss of a predicted start codon. The gene contains a single exon, encodes a protein length of 550 amino acids (aa) and is predicted to contain a fungal-specific transcription factor domain and Zn(2)-C6 fungal-type DNA-binding domain (Table 4). The orthologue in the 3D1 genome is *ZT3D1_G3567*. It encodes a longer protein (722 aa) and has two exons, the first exon’s product encodes a Zn(2)-C6 fungal-type DNA-binding domain and the second exon’s product is the same as that of *ZT3D7_G3622*. The predicted protein in 3D1 is the same as the orthologue in IPO323 (*Mycgr3T108446*). Although we do have reads mapping to this region, and when grown in vitro on MM this gene is downregulated in 3D1 compared to 3D7 (Table S8), our RNA sequence data does not clearly support gene models in either 3D1 or 3D7, there are no clear exon/intron boundaries. This suggests that the annotation of this gene may require manual curation.

*ZT3D7_G3635* has a high impact SNP variant (Chr 3:681906) that is predicted to cause a frameshift. The gene has 6 exons in 3D7 and encodes a protein length of 1078 aa. Based on INTERPROSCAN and Ensemble Fungi, the protein has multiple predicted domains; the largest is a protein kinase domain. The protein also contains a Serine/threonine-protein kinase active site and the molecular function of the first GO term (GO:0004674) is protein serine/threonine kinase activity – suggesting that this gene is likely to be involved in osmotic stress response (Hohmann 2002). The RNA sequence data supports the gene model. The orthologue in 3D1 is *ZT3D1_G3580*, which has 3 exons and multiple predicted domains that do not differ from those predicted for *ZT3D7_G3635*. The gene is up-regulated in vitro on YSB and down-regulated in planta at 28d in 3D1 compared to 3D7 (Table S8).

*ZT3D7_G3637* has one exon, encodes a protein length of 1285 aa, and includes multiple predicted domains including a DEAD/DEAH box helicase, which is involved in plant responses to osmotic stress (Nguyen *et al*. 2018). The orthologue in 3D1 (*ZT3D1_G3583*) encodes a similar protein as *ZT3D7_G3637* – has the same length and has the same predicted domains. There are no high impact SNPs between the parents, but several moderate and low impact variants were identified (Table S*). The RNA sequence data supports the annotation, and the gene is down regulated in 3D1 in both YSB and MM (Table S8).

*ZT3D7_G3639* has 4 exons in 3D7. The gene sequence similarity between 3D7 and 3D1 is 98.66% and there is an 8 bp intronic deletion in 3D1. This gene belongs to the Major Facilitator Superfamily (MFS) and encodes a protein length of 512 aa. According to the Phobius analysis, the gene contains 12 transmembrane segments (Table S8). Proteins with transmembrane activity and transmembrane segments are likely to play an important role in sensing and regulating osmotic stress (Hohmann 2002). The orthologue in 3D1 is *ZT3D1_G3580*, which has the same number of exons, encodes the same protein length and has the same predicted function. The RNA sequence data supports the gene model, and the differential expression analysis found the gene is downregulated in planta at 12 and 14 days in 3D1 compared to 3D7 (Table S8).

*ZT3D7_G3655* has one exon and encodes a protein length of 848 aa. The gene contains 10 transmembrane segments (Table S8) and encodes a voltage-gated chloride channel domain – thus it is likely to be involved in osmotic stress responses. The gene sequence between the parents is almost identical, but there is one moderate effect SNP within the gene (Table S9). When grown in vitro on MM this gene is upregulated in 3D1 compared to 3D7. The RNA sequence data supports the gene model annotation.

*ZT3D7_G3658* has four exons and encodes a protein length of 614 aa. Its orthologue in 3D1 (*ZT3D1_G3604*) has three exons and encodes 572 aa. The gene encodes a vacuolar protein sorting-associated protein 17, Vps17. Vacuoles play an important role in osmotic stress responses: their fission and fusion enable cells to maintain turgor pressure and regulate cell volume (Richards *et al*. 2010). Comparison of the 3D1 gene sequence to the 3D7 genome using BLAST revealed that the 3D7 gene sequence was identical, but the 3D1 orthologue annotation is shorter and has only three exons. Our RNA sequence data supports the 3D1 annotation with only 3 exons, as there were no RNA sequence reads mapping to exon 1 of the *ZT3D7_G36580*. This gene was downregulated in vitro in both MM and YSB in 3D1 compared to 3D7.

## 4. Discussion

We were able to map major effect loci affecting colony growth and melanisation in the presence of potassium chloride in both crosses. We focussed on QTL that were unique to the KCl environment to identify candidate genes that are specific to osmotic stress – as opposed to finding genes that may be more generally related to intrinsic growth or melanin production. A QTL on Chr 3 in the 3D7×3D1 cross was the most promising for identifying genes specifically associated with KCl stress. We highlighted the six most promising candidate genes in this QTL based on their predicted encoded functions, sequence similarity and transcriptional profiles. We also identified several QTL shared across environments and confirmed that several previously identified QTL, which explained variation in growth and melanisation across different environmental stressors (Table 2 and 3, e.g. 3D7×3D1 Chr 11 and 1A5×1E4 Chr 8), also explained trait variation in the presence of osmotic stress.

### Effect of KCl on colonies

In the presence of potassium chloride at 0.75M, colonies were typically smaller and less melanised compared to the control environment (Figure 1, Table S2). A reduction in colony radius in the KCl environment supports our hypothesis that this environment was stressful to *Z. tritici*. Our results suggest that growth rate in the 1A5×1E4 cross was less affected by KCl-induced osmotic stress, as mean growth rate was not reduced in the KCl environment, compared to the 3D7×3D1 cross that exhibited a significantly reduced growth rate (Table S2). However, we note that the overall difference in growth rate in the 3D7×3D1 progeny between the control and KCl environments was relatively small (∼6% reduction in growth, Table S2).

When experiencing osmotic stress, an increase in melanin is thought to protect cells by strengthening the fungal cell wall, but a reduced metabolic capacity under stress may limit the ability of a cell to produce melanin. The mean colony grey values were higher, indicative of lower levels of melanin, for strains grown in the KCl environment (Table S2). This suggests that the *Z. tritici* response to osmotic stress did not involve an increased production of melanin to protect the cells as was observed in halotolerant species (Kogej *et al*. 2007). We did observe however, that isolates that were more melanised in the benign control environment (lower Grey Value) were more tolerant to salt stress (Figure S2) – their growth rates were higher and their reductions in melanisation were lower compared to isolates that were less melanised in the control environment (Table S3, S4). This suggests that strains having a greater intrinsic capacity to become melanised are more tolerant to salt stress.

### Shared QTL

Some QTL that explained variation in growth and melanisation in the stressful environment overlapped with QTL for trait variation in the control environment. Prominent examples include the Chr 11 QTL No. 11 in the 3D7×3D1 cross and the Chr 8 QTL No. 10 in the 1A5×1E4 cross. These genomic regions may harbour genes that affect intrinsic growth of colonies under a wide range of environments and/or genes with pleiotropic effects. Many of the QTL shared across environments were identified in earlier experiments and discussed previously (Lendenmann *et al*. 2015; Zhong *et al*. 2021). The QTL No. 11 in the 3D7×3D1 was shown to be due to differential expression of the *Zmr1* gene encoding a transcription factor that regulates synthesis of DHN melanin in *Z. tritici* (Krishnan et al. 2018). *Zmr1* was shown to have pleiotropic effects on melanin production, colony growth rates and fungicide resistance (Krishnan et al. 2018).

### QTL uniquely associated with the KCl environment

This QTL analysis enabled us to identity large effect loci that explain variation in traits under KCl stress. Among the QTL uniquely associated with KCl stress, most had moderate LODs with each QTL explaining only a small proportion of the phenotypic variance (3D7×3D1 maximum = 0.10, 1A5×1E4 maximum = 0.11). The exception was QTL No. 3 in the 3D7×3D1 cross, which explained up to 33% of the phenotypic variance. This QTL on chromosome 3 was associated with differences in colony radius at 8 dpi and colony radius KCl tolerance at 8 and 12 dpi. We targeted this QTL for further investigation, seeking to identify genes specifically associated with KCl tolerance. It is interesting to note that no LOD peak was detected for colony radius at 12 dpi, even though colony radius at 12 dpi was correlated with colony radius at 8 dpi (Table S4, correlation = 0.58, p<0.001). As growth rate is based on the change in colony radius between 8 and 12 dpi, we might expect growth rate in KCl to share the QTL for colony radius at 8 dpi. To better understand why growth rate does not share this QTL, we examined the trait correlations. Growth rate is highly correlated with colony radius at 12 dpi (0.93) and less strongly correlated with colony radius at 8 dpi (0.26) (Table S4). There is less variation in colony radius at 8 dpi, thus colony radius at 12 dpi has a greater impact on the growth rate. The QTL No. 3 also explained variation in colony radius KCl tolerance at 8 and 12 dpi, although the association with this trait at 12 dpi explained relatively little phenotypic variation (0.08). Thus, the QTL peak is more strongly associated with colony radius and colony radius KCl tolerance at an earlier stage of colony growth. A similar pattern was observed in the chromosome 11 QTL associated with melanisation (Lendenmann *et al*. 2014; Krishnan *et al*. 2018), with the QTL LOD score very high in the younger colonies and decreasing as colonies aged. After cloning and characterizing the *Zmr1* gene and its functions, this pattern could be explained based on the finding that the difference in *Zmr1* expression (and the resulting difference in melanisation) was significantly higher in younger colonies than in older colonies (Krishnan *et al*. 2018). We hypothesize that QTL no. 3 associated with KCl tolerance is underpinned by a similar mechanism that is specific to the earlier stage of colony growth.

### Candidate genes in the QTL for colony radius and colony radius KCl tolerance at 8 dpi in the 3D7×3D1 cross

Among the 36 genes found in the interval spanning this QTL No. 3, we consider six genes to be the most promising candidates to be involved in sensing and responding to salt stress and responsible for this QTL. Below we consider each in turn in the order of their relative positions on chromosome 3.

The first candidate gene, *ZT3D7_G3622*, is predicted to encode a fungal-specific transcription factor. The orthologue in the IPO323 reference genome (assembly MYCGR v2.0) is *Mycgr3T108446* and it has 21 paralogs. Transcription factors (TFs) regulate gene expression and are important regulators of molecular responses to environmental stimuli (e.g. Hohmann, 2002; Shi et al., 2021), making this gene a good candidate. The *Zmr1* gene identified in this same cross and shown to be responsible for differences in melanisation, with greater effects in younger colonies, is a transcription factor (Krishnan et al. 2018). The SNP variant in 3D1 (Chr 3:598092) is predicted to cause a loss of the start codon that is likely to cause a loss of function. In *F. graminearum,* mutants lacking a similar TF (called *Fss1*) were sensitive to high sodium and lithium, but they were not sensitive to elevated potassium (Son *et al*. 2015).

The second candidate, *ZT3D7_G3635* is predicted to encode a serine/threonine-protein kinase active site, which is known to regulate MAP kinases (Hohmann 2002; Li *et al*. 2019). A SNP variant found in the 3D1 allele is predicted to cause a frameshift that is likely to cause a loss of function. The ortholog in the IPO323 genome is *Mycgr3G56270* and it has one paralog in the reference genome. A serine/threonine kinase from *Aspergillus flavus*, *AflSte20*, was recently functionally validated using a knockout mutant. This study showed that the knockout had higher sensitivity to osmotic stress compared to the wild type (Li *et al*. 2019), confirming the role of this serine/threonine-protein kinase in osmotic stress response in a different fungal plant pathogen.

The third candidate, *ZT3D7_G3637* is predicted to encode a DEAD/DEAH box helicase domain. The reference genome orthologue is *Mycgr3G99415* and it has 6 paralogs. The DEAD – box helicases are the largest RNA helicase (RH) subfamily. In plants they play multiple functional roles and are important in stress responses (Pandey *et al*. 2020). For example, in *Arabidopsis thaliana AtRH17* is involved in salt-stress tolerance (Nguyen *et al*. 2018), in rice *OsABP* is upregulated in response to salt (Macovei *et al*. 2012) and in barley HvD1 accumulated in salt- and cold-stressed plants (Nakamura *et al*. 2004). In wheat, the DEAD-box helicase *TaRH1* that was first identified as a component of the defence response to stripe rust (*Puccinia striiformis f*. sp. *tritici*) was also found to be involved in plant response to multiple stressors, including salt stress (Zhang *et al*. 2014). SNP variation found between the parents was predicted to have only moderate effects, no high impact SNPs were detected. We chose this gene as a promising candidate based mainly on its known associations with salt stress in plants and the significant differences in expression found between 3D1 and 3D7 while growing in vitro.

A gene encoding a Major Facilitator Superfamily (MFS) domain, *ZT3D7_G3639* is the fourth candidate we highlight. The reference genome orthologue is *Mycgr3G85019* and it has 58 paralogs. MFS transporters facilitate the movement of small solutes across cell membranes in response to chemiosmotic gradients – making them likely to play an important role in sensing and regulating osmotic stress (Hohmann 2002). We chose this gene as a promising candidate based mainly on its known functions and associations with multiple stressors including fungicides (Dos Santos *et al*. 2014; Omrane *et al*. 2015).

The fifth highlighted gene is *ZT3D7_G3655* that encodes a voltage gated chloride channel domain and membrane transport domains. The reference genome orthologue is *Mycgr3T20761* and there are three paralogues in the IPO323 reference genome. Chloride channels are important in animal cell volume regulation (Jentsch *et al*. 2002), fungal oxidative stress response (Oddon *et al*. 2007), fungal pathogen virulence (Zhu and Williamson 2003; Cañero and Roncero 2008) and plant immunity (Guo *et al*. 2014). The role of chloride channels in response to salt stress is well established in animals (Jentsch *et al*. 2002), and to a lesser degree in plants (Barbier-Brygoo *et al*. 2011), but less is known from fungi. For one of the better-known families of chloride channels, the CLC family, a single gene (*GEF1*) has been identified in yeast. In two pathogenic fungi (*Cryptococcus neoformans* and *Fusarium oxysporum*) – the GEF1 homologue has been shown to influence virulence (Zhu and Williamson 2003; Cañero and Roncero 2008). The gene sequence between the parents is highly conserved, however gene expression differences were observed in minimal media, so differences may exist in the regulatory sequence.

The final candidate we highlight is *ZT3D7_G3658*, which encodes a Vacuolar associated protein 17, *Vps17*. The orthologous gene in the reference genome is *Mycgr3T108464* and there are six paralogues. Comparing the DNA sequence of this gene against the 3D1 genome revealed an identical sequence, but the annotation of the gene in 3D7 and 3D1 differs – the 3D1 orthologue protein length is shorter and the gene has one fewer exons. Our RNA sequence data suggests that the *ZT3D7_G3658* annotation may be incorrect - there were no reads mapping to the first, small exon of this gene. This suggests that the annotation of this gene may need manual curation before conducting further analyses. The gene was downregulated in YSB and MM in 3D1 compared to 3D7, demonstrating that transcription of this gene varies between the parents. Vacuoles play important roles in both osmotic stress responses (Richards *et al*. 2010) and virulence (Li *et al*. 2021). Under salt stress the vacuoles can undergo fusion and fission – changing in size and number (Richards *et al*. 2010). *Vsp17* knockouts in *S. cerevisiae* had many small vacuoles but were resistant to osmotic stress and could sporulate (Köhrer and Emr 1993). We are unaware of any work that has investigated the role of *Vsp17* in salt-stress responses of filamentous fungi.

### Enrichment analysis of QTL unique to the KCl environment

We investigated if genes within QTL unique to the KCl environment were functionally enriched. We considered each cross separately and considered colony size-related traits and melanin-related traits separately (Table 2, 3: QTL No.^S or M^). No KEGG pathway enrichment was detected. This may reflect the relatively low number of genes that were assigned a K number (3D7 34%, 1A5 49%). The GO enrichment analysis provided some interesting results. For QTL associated with colony size related traits, two notable GO annotations were enriched. The first was cell wall (GO:0005618) which was enriched in QTLs in the 1A5×1E4 cross. Cell wall remodelling is an important response to osmotic stress and this remodelling is mediated by the Hog1 and MAPK pathways (as described earlier). The cell wall GO is associated with four genes in the QTL intervals: ZT1A5_G201, ZT1A5_G1063, ZT1A5_G1074, and ZT1A5_G1774. A common annotation feature of the first two genes is they contain a glycoside hydrolase family 16, which is involved in degradation and remodelling of the cell wall. In rice blast (*M. oryzae*), glycoside hydrolases II homolog (MoGls2) mutants showed defects in cell wall integrity, were less sensitive to salt stress and had reduced virulence (Li *et al*. 2016). The other two genes with this GO term (ZT1A5_G1074, ZT1A5_G1774) contain yeast PIR protein repeat. A number of yeast cell wall glycoproteins are characterised by the presence of internal repeats (PIR, tandem repeats of 18 to 19 amino acid residues) and these are important in yeast response to multiple environmental stressors and antifungal agents (Mazáň *et al*. 2008). The second notable GO enrichment was for GO terms related to DNA repair (GO:0006281: nucleotide-excision repair, double-strand break repair via nonhomologous end joining). DNA repair genes can be upregulated in the presence of a variety of environmental stressors and are widely considered an important response to stress (Hohmann 2002). Six genes have these GO annotations (ZT3D7_G5293, ZT3D7_G3673, ZT3D7_G3591, ZT3D7_G5301, ZT3D7_G5305, ZT3D7_G5361). Three of these genes (ZT3D7_G5293, ZT3D7_G3673, ZT3D7_G53601) have multiple GO annotations, of which three were also enriched: DNA binding (GO:0043564), damaged DNA binding (GO:0003684), and telomeric DNA binding (GO:0042162). Another three genes with enriched GO annotations were in or adjacent to the large LOD QTL on chromosome 3 (Figure 4, Table 4) in the 3D7×3D1 cross (ZT3D7_G3648, ZT3D7_G3652, ZT3D7_G3659, Table S7). Two have predicted GTPase activity (GO:0003924, ZT3D7_G3648 and ZT3D7_G3659). GTPase are involved in many functions in fungi, including during plant-fungal interactions. A well-studied GTPase family, *Ras*, has been shown to be involved in response to osmotic stress in *Beauveria bassiana* and *Botrytis cinerea* (Dautt-Castro *et al*. 2021). Fungal treatments targeting Ras proteins are being tested to control human pathogens with promising results (LeBlanc *et al*. 2020). The other gene in the interval, ZT3D7_G3652 has predicted superoxide dismutase activity (GO:0004784). Genes in this family are known to play an important role in defending cells against reactive oxygen species, and in the stripe rust fungus (*Puccinia striiformis* f. sp. *Tritici*) this can help guard against plant defences and facilitate infection (Zheng *et al*. 2020).

There was also a notable GO term enriched for melanin-related QTL: glutathione biosynthetic process (GO:0006750). This GO term was associated with three genes within QTLs: ZT3D7_G1409, ZT3D7_G1830, ZT3D7_G8325. Glutathione is a metabolite that plays a major role in response to stress, morphogenesis, and virulence in fungi (Wangsanut and Pongpom 2022). In *Cryptococcus neoformans,* a glutathione gene mutant (gsh2Δ) had increased sensitivity to salt (Berndt and Lillig 2017), and *Schizosaccharomyces pombe* grown in vitro under multiple stressors, including osmotic stress, had increased expression of the glutathione reductase gene (*pgr1*) (Lee *et al*. 1997). Our results suggest that glutathione may also be important in osmotic stress response in *Z. tritici*. All the genes highlighted in our GO enrichment analysis may be useful targets for further investigation to understand the molecular response of *Z. tritici* to salt stress, but may also provide interesting targets for fungicide development.

### QTL reproducibility

This dataset allowed us to explore the repeatability of previously identified QTL from other published datasets. One earlier dataset used the same mapping populations but different experimental conditions and treatments, with a different reference genome and different genetic markers and map information (Lendenmann et al., 2015, 2014). A second earlier dataset used the same isolates and experimental conditions, but different treatments and different marker and map information (Zhong *et al*. 2021). Comparisons across these three data sets yielded several consistent QTL. Our analyses confirmed that a region on chromosome 11 in the 3D7×3D1 cross (Table 2, QTL No. 11) is associated with variation in melanisation and growth in multiple environments (e.g. under fungicide stress in Lendenmann et al., 2015; under oxidative stress in Zhong et al., 2021). This QTL contains the *Zmr1* transcription factor that was shown to regulate melanin production (Krishnan *et al*. 2018). We hypothesize that *Zmr1* also explains the variation we observed under salt-stress. A QTL associated with multiple growth and melanin traits in the 1A5×1E4 cross on chromosome 8 (Table 3, QTL No. 13) overlaps with a previously identified QTL on chromosome 8 that was associated with colony growth and tolerance to oxidative stress (Zhong *et al*. 2021). These findings illustrate that many of our QTL are reproducible across experiments and increases our confidence that we have identified biologically meaningful genomic regions using our automated image analyses.

## Authorship contribution statement

Jessica Stapley: Formal analysis, Data curation, Writing - Original Draft, Writing - Review & Editing, Resources, Data curation. Bruce A. McDonald: Conceptualization, Methodology, Writing - Review & Editing, Supervision, Project administration, Funding acquisition.

## Data availability

Phenotypic and SNP data are available at ETH Data repository (DOI: 10.3929/ethz-b-000550424). All code and details of the analysis are available at (https://github.com/jessstapley/QTL-mapping-Z.-tritici).

## Acknowledgments

We thank Ziming Zhong, Tiziana Valeria Vonlanthen, Bethan Turnbull, Susanne Dora, Sarah Furler, Alexandra Waltenspühl, and Jasmin Wiedmer for collecting the phenotypic data and Javier Palma-Guerrero for supervision. We thank Marcello Zala for assistance in the laboratory. This research was supported by the Swiss National Science Foundation (31003A_155955 granted to BAM).

## Supplementary Figures

Figure S1. Correlation of growth rate and melanisation rate in control and KCl environment.

Figure S2. Correlation of mean radius and mean grey value ar 12 dpi in control and KCl environment.

Figure S3. LOD plots from interval mapping for all traits in the 3D7×3D1 cross.

Figure S4. LOD plots from interval mapping for all traits in the 1A5×1E4 cross.

## Supplementary Tables

Table S1. Linkage map summary

Table S2. Summary of phenotypic data

Table S3. Trait correlations in 3D7×3D1 cross.

Table S4. Trait correlations in 1A5×1E4 cross.

Table S5. Results of the GO enrichment analysis for 3D7×3D1 cross.

Table S6. Results of the GO enrichment analysis for 1A5×1E4 cross.

Table S7. GO enrichment genes.

Table S8. QTL interval on Chr 3 (QTL#3) in 3D7×3D1 cross in detail.

Table S9. Results of the SNP effect analysis for the QTL interval on Chr 3 (QTL #3)

