## Supplementary figures and images for "Quantitative trait locus mapping of osmotic stress response in the fungal wheat pathogen *Zymoseptoria tritici*"

### Figure S1

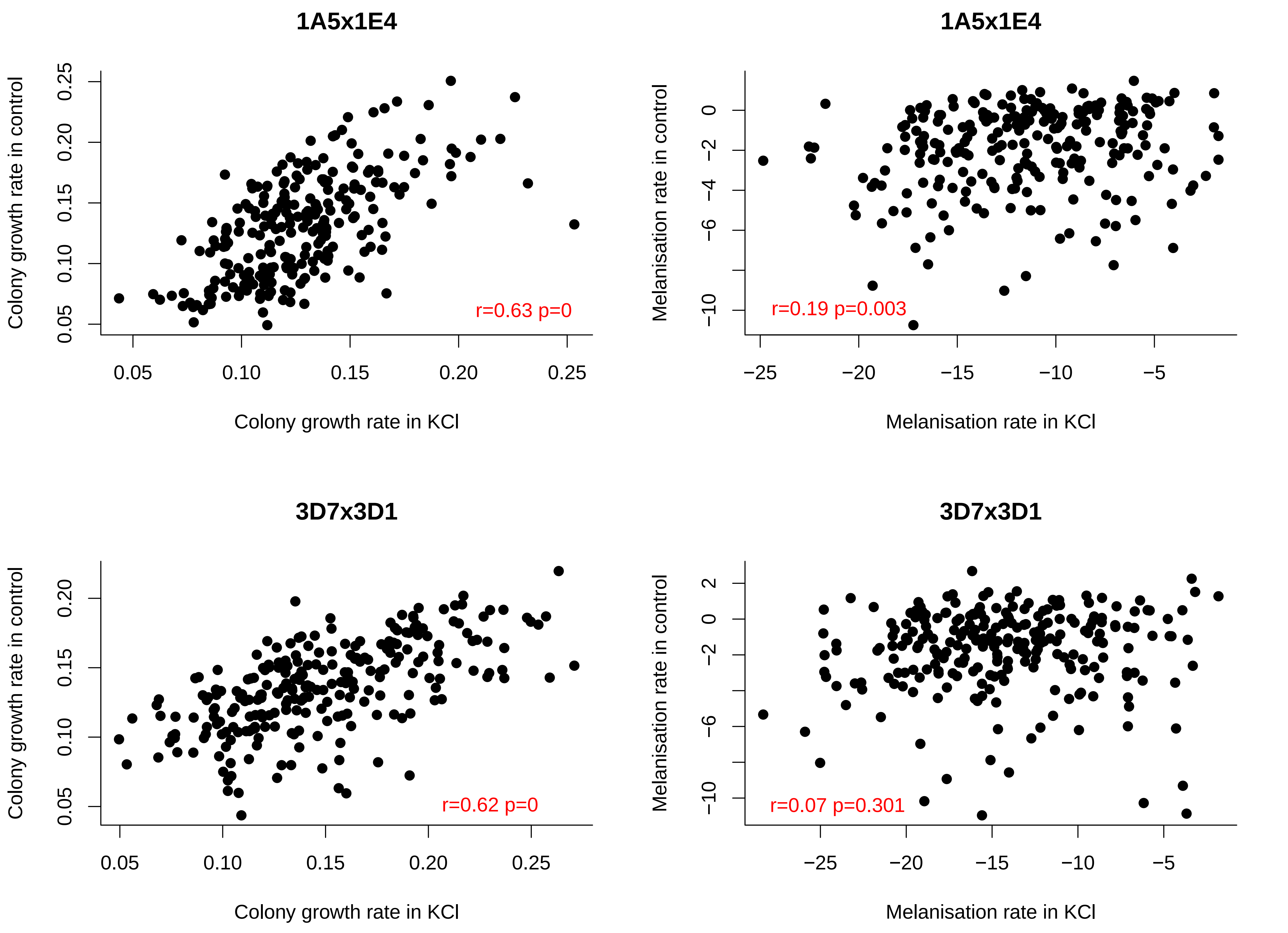

### Figure S2

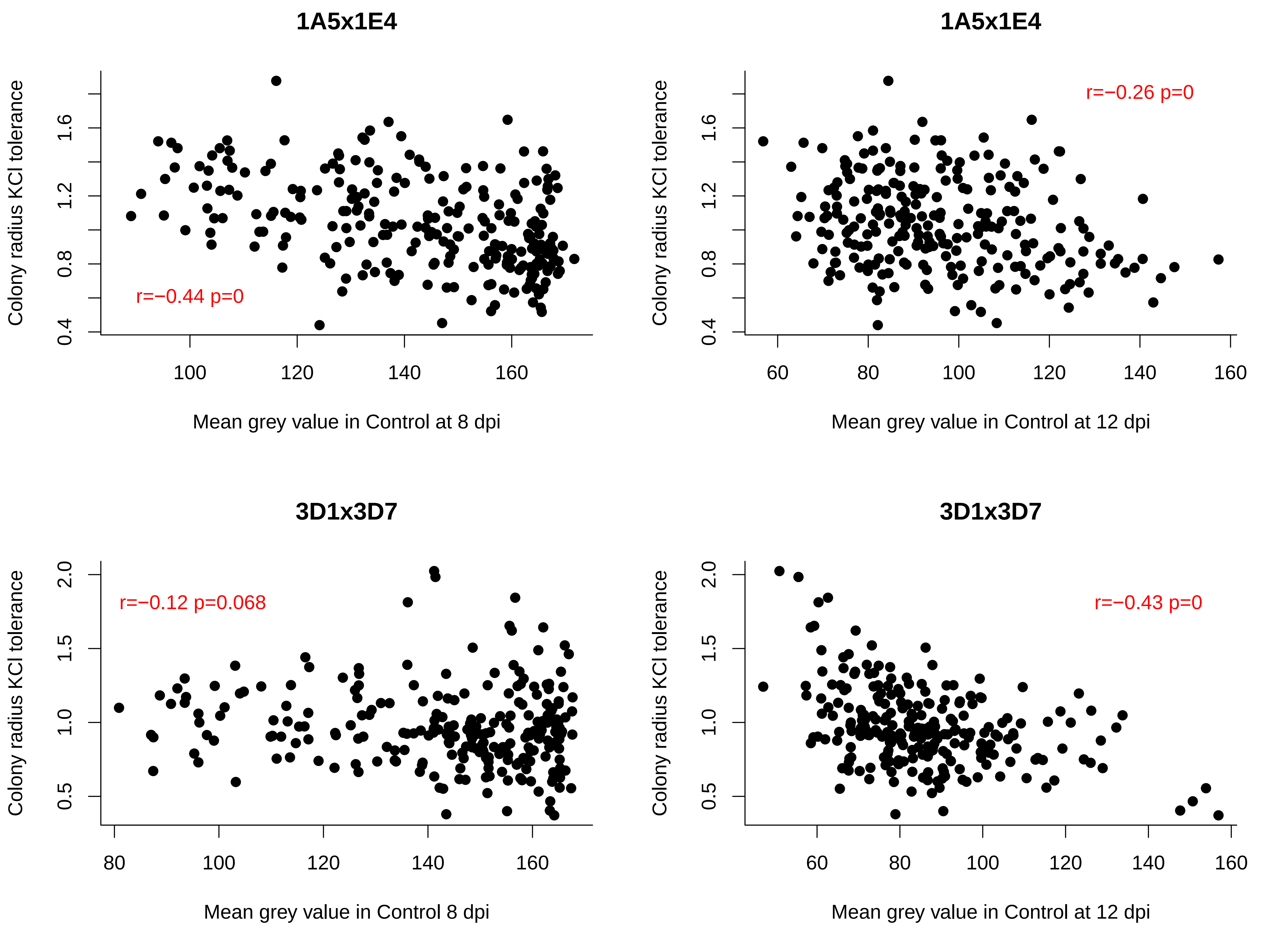
