## Supplementary material for "Quantitative trait locus mapping of osmotic stress response in the fungal wheat pathogen *Zymoseptoria tritici*": Figure S3

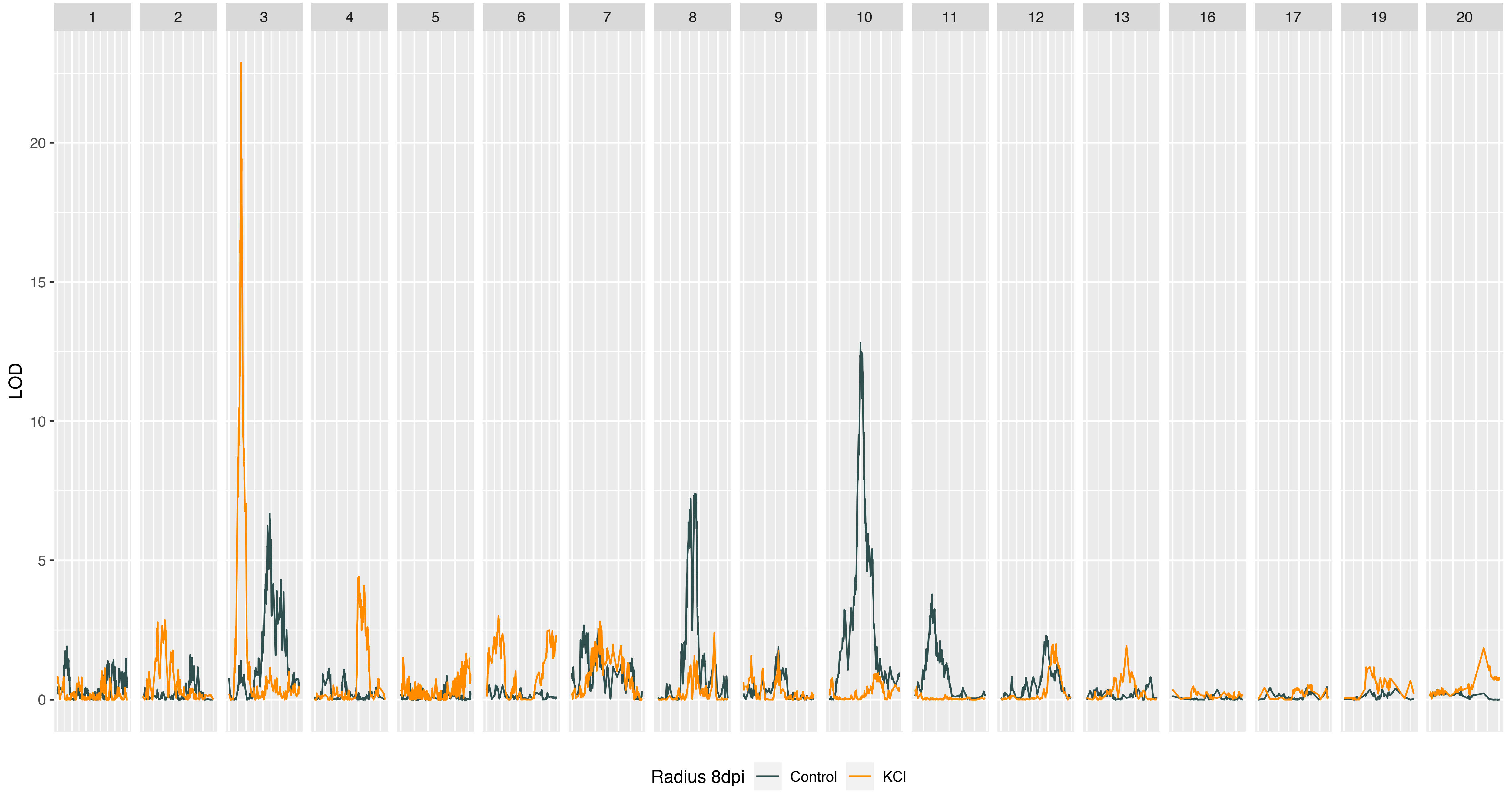
1
2
3
4
5
6
7
8
9
10
11
12
13
16
17
19
20
LOD
20
15
10
5
0
Radius 8dpi
Control
KCl

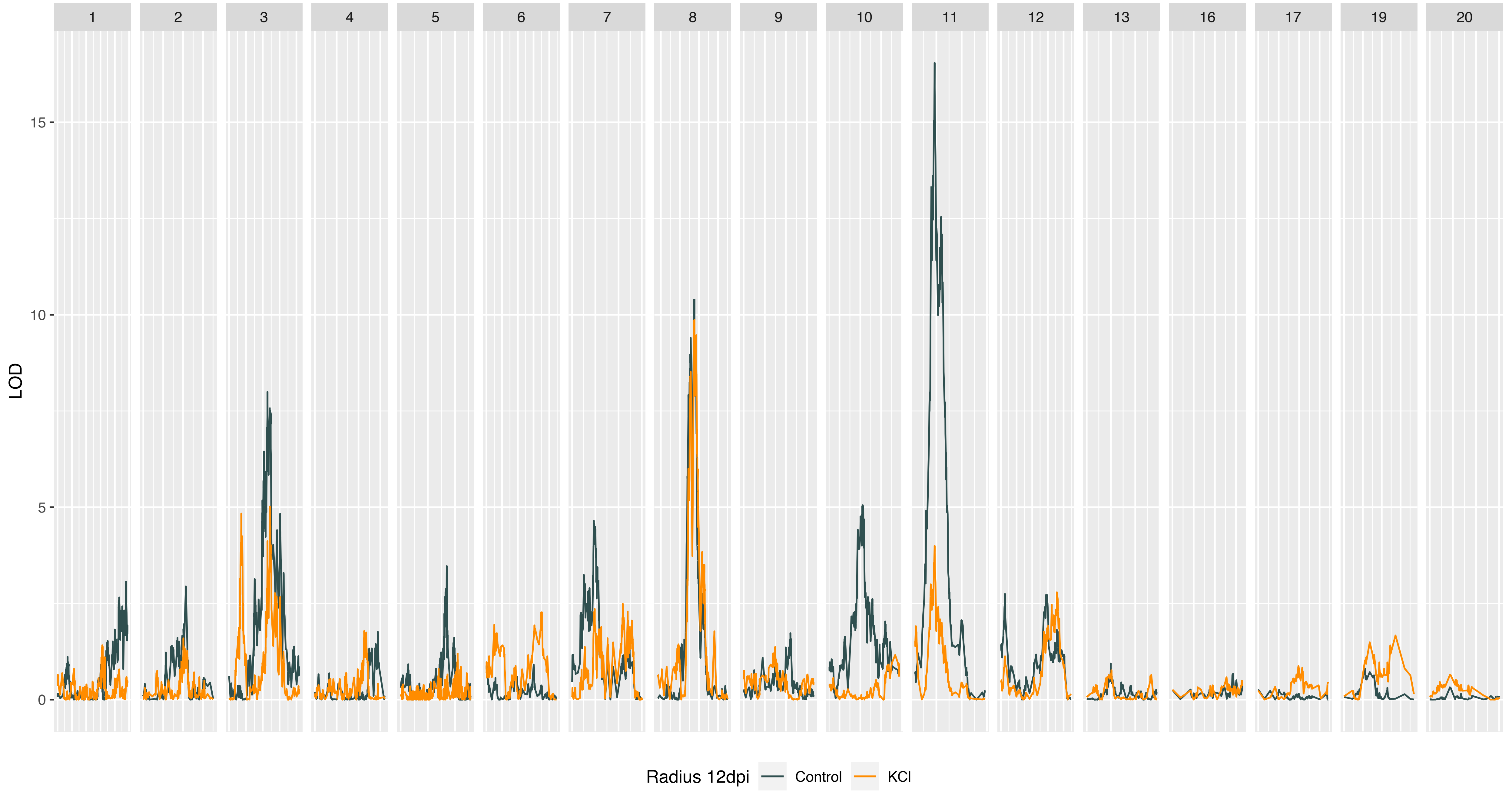

1
2
3
4
5
6
7
8
9
10
11
12
13
16
17
19
20
LOD
15
10
5
0
Radius 12dpi
Control
KCl

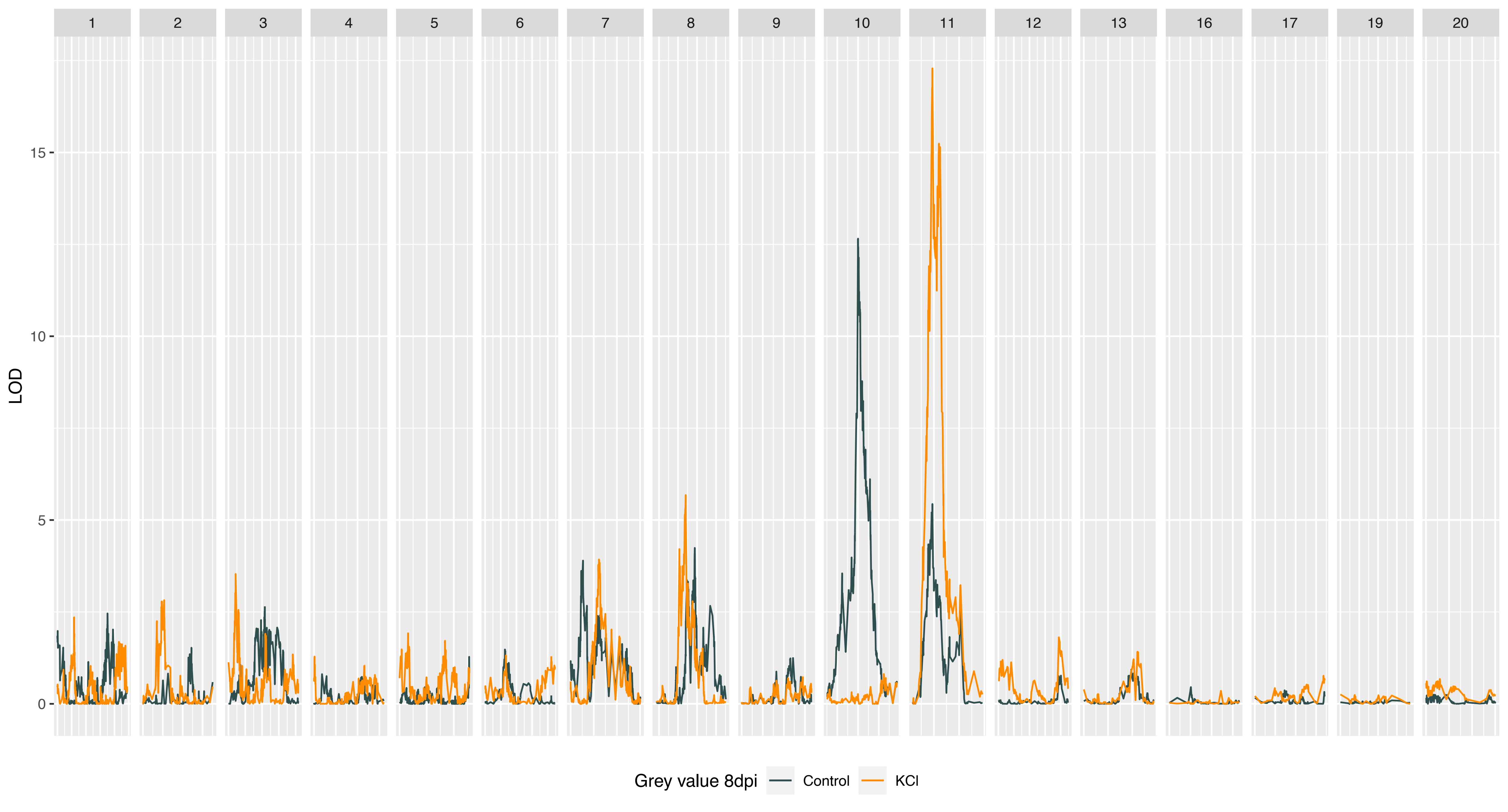
1
2
3
4
5
6
7
8
9
10
11
12
13
16
17
19
20
LOD
15
10
5
0
Grey value 8dpi
Control
KCl

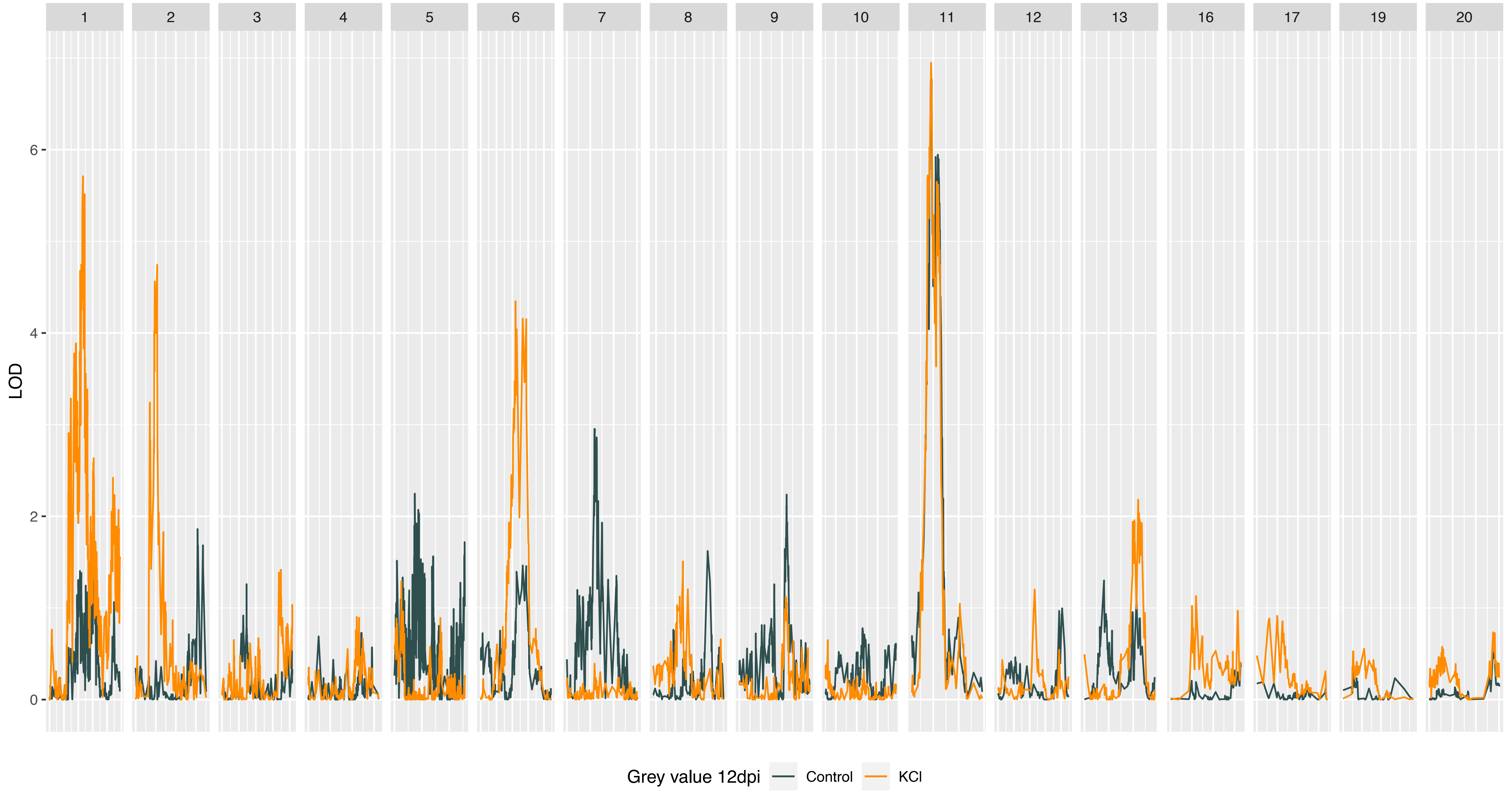

1
2
3
4
5
6
7
8
9
10
11
12
13
16
17
19
20
LOD
6
4
2
0
Grey value 12dpi
Control
KCl

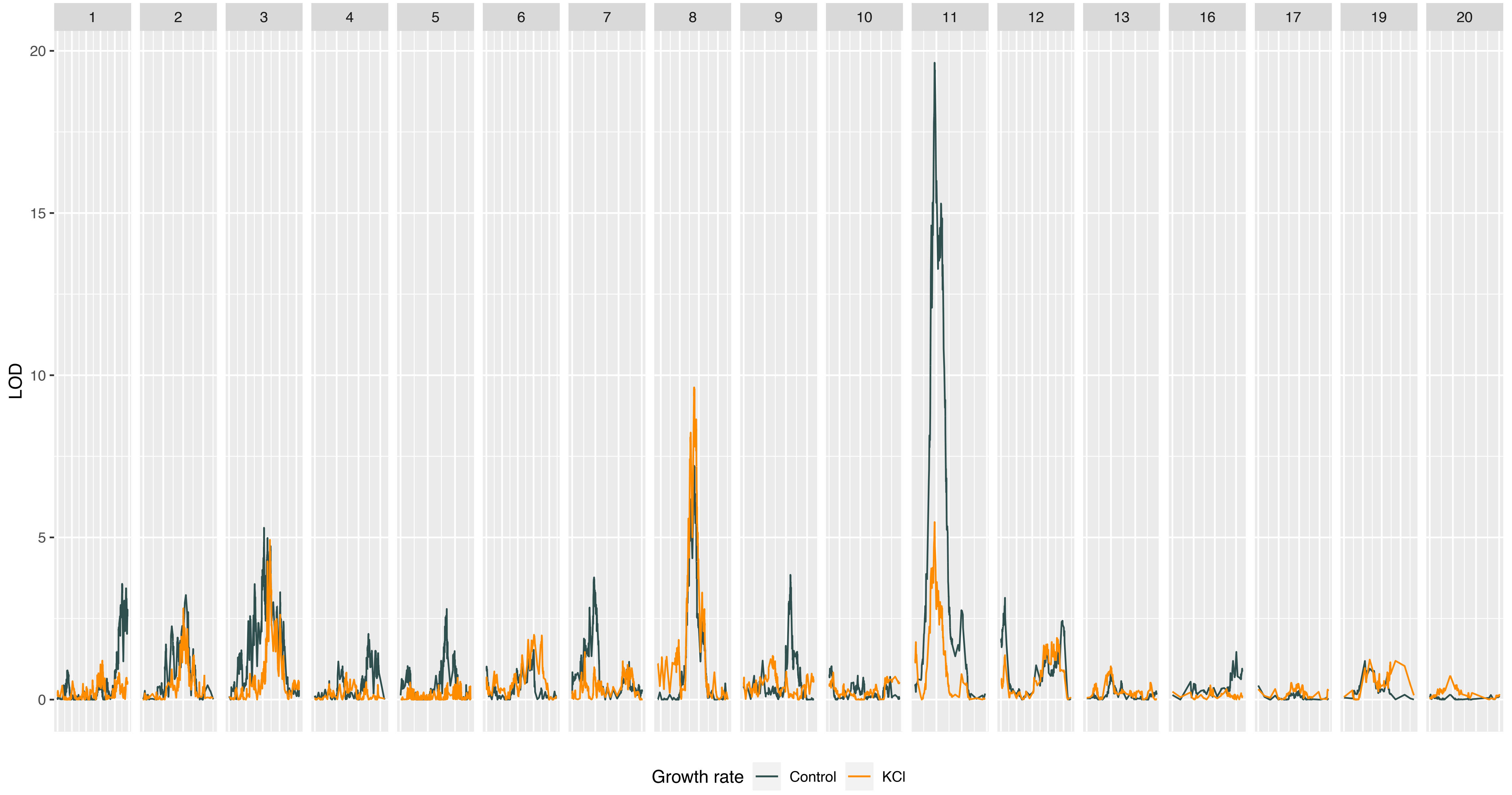
1
2
3
4
5
6
7
8
9
10
11
12
13
16
17
19
20
LOD
20
15
10
5
0
Growth rate
Control
KCl

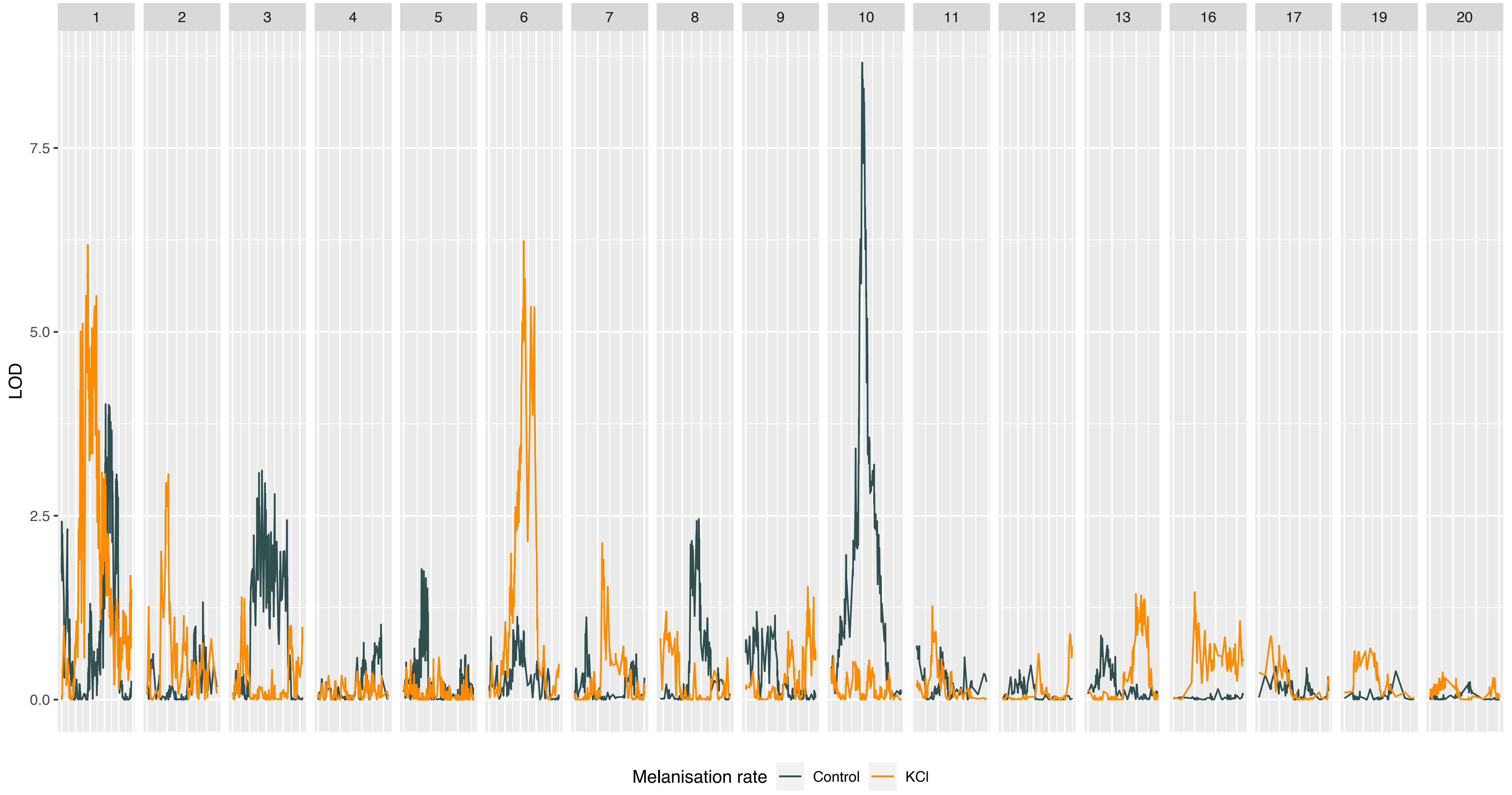

1
2
3
4
5
6
7
8
9
10
11
12
13
16
17
19
20
LOD
7.5
5.0
2.5
0.0
Melanisation rate
Control
KCl

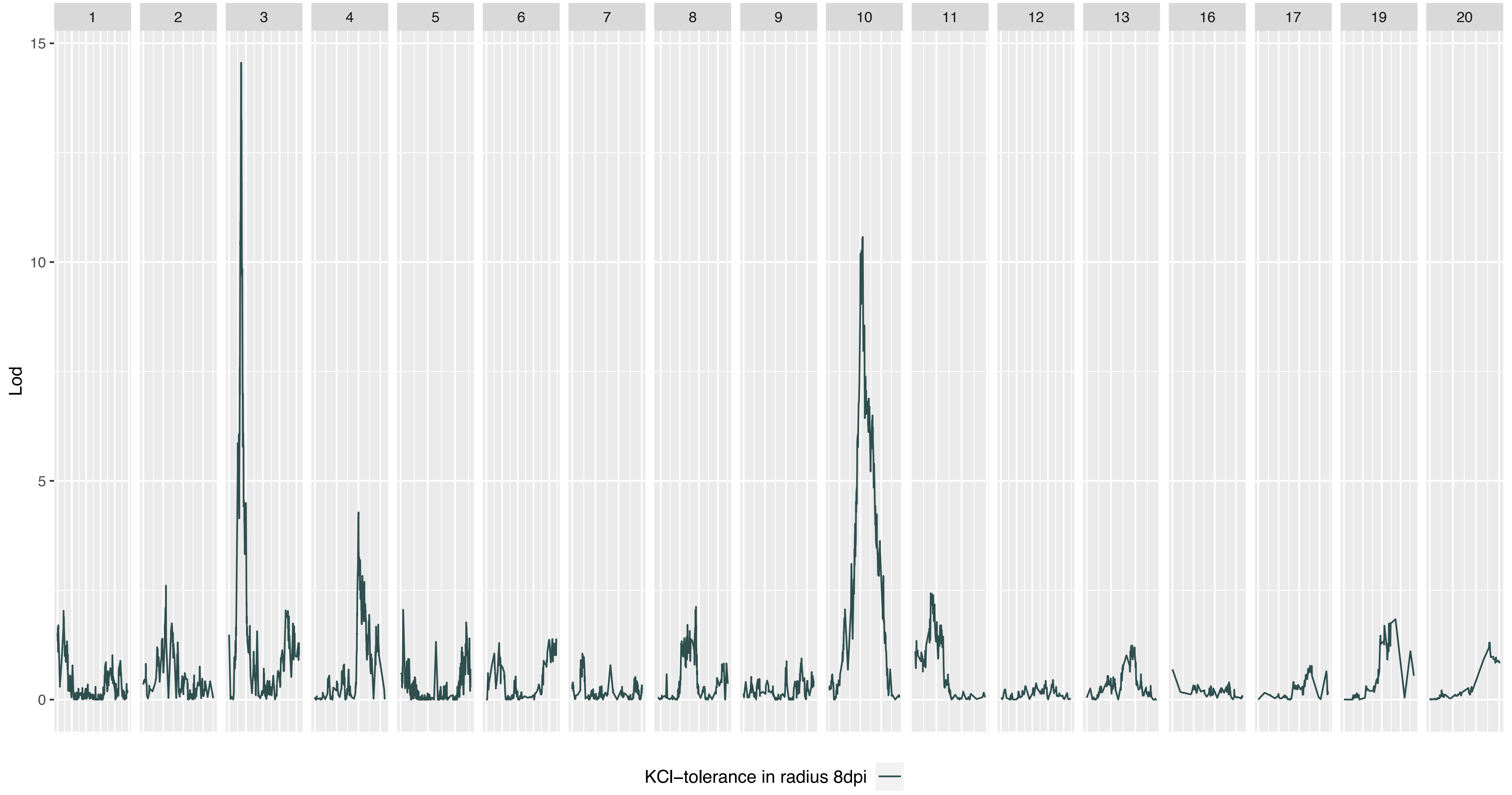

1
2
3
4
5
6
7
8
9
10
11
12
13
16
17
19
20
15
10
5
0
Lod
KCl–tolerance in radius 8dpi

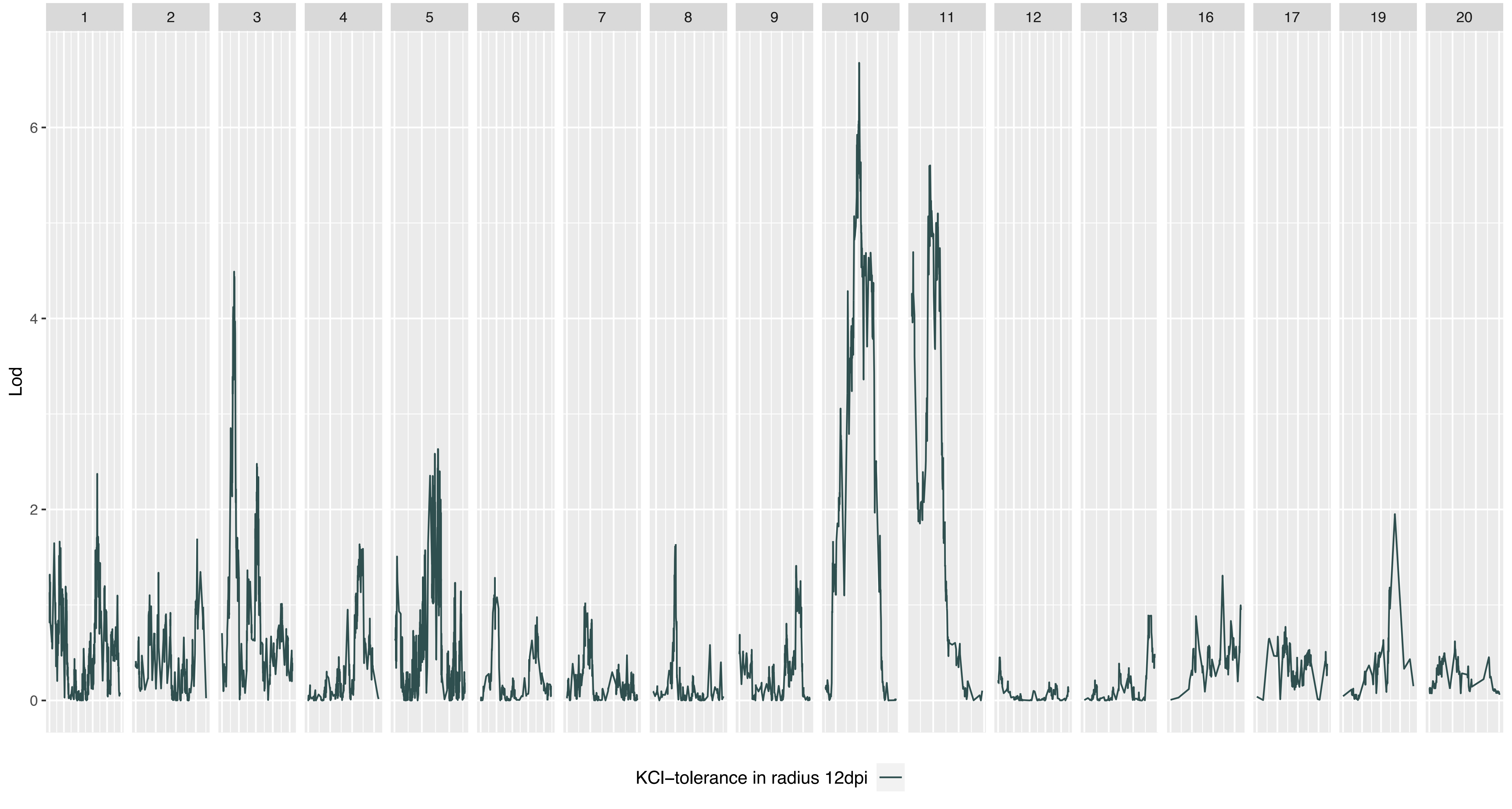
1
2
3
4
5
6
7
8
9
10
11
12
13
16
17
19
20
Lod
0
2
4
6
KCl–tolerance in radius 12dpi

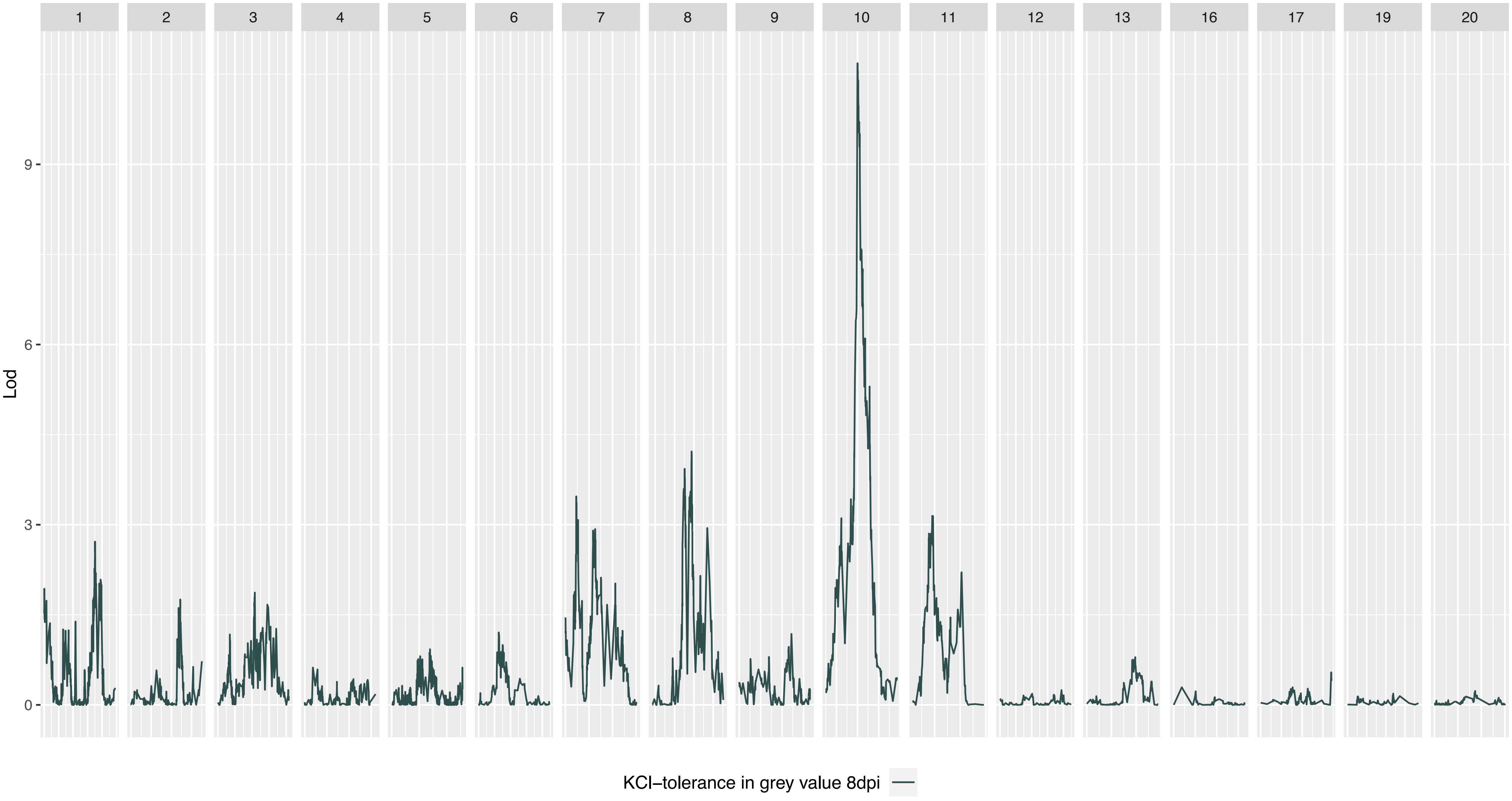

1
2
3
4
5
6
7
8
9
10
11
12
13
16
17
19
20
Lod
0
3
6
9
KCl−tolerance in grey value 8dpi

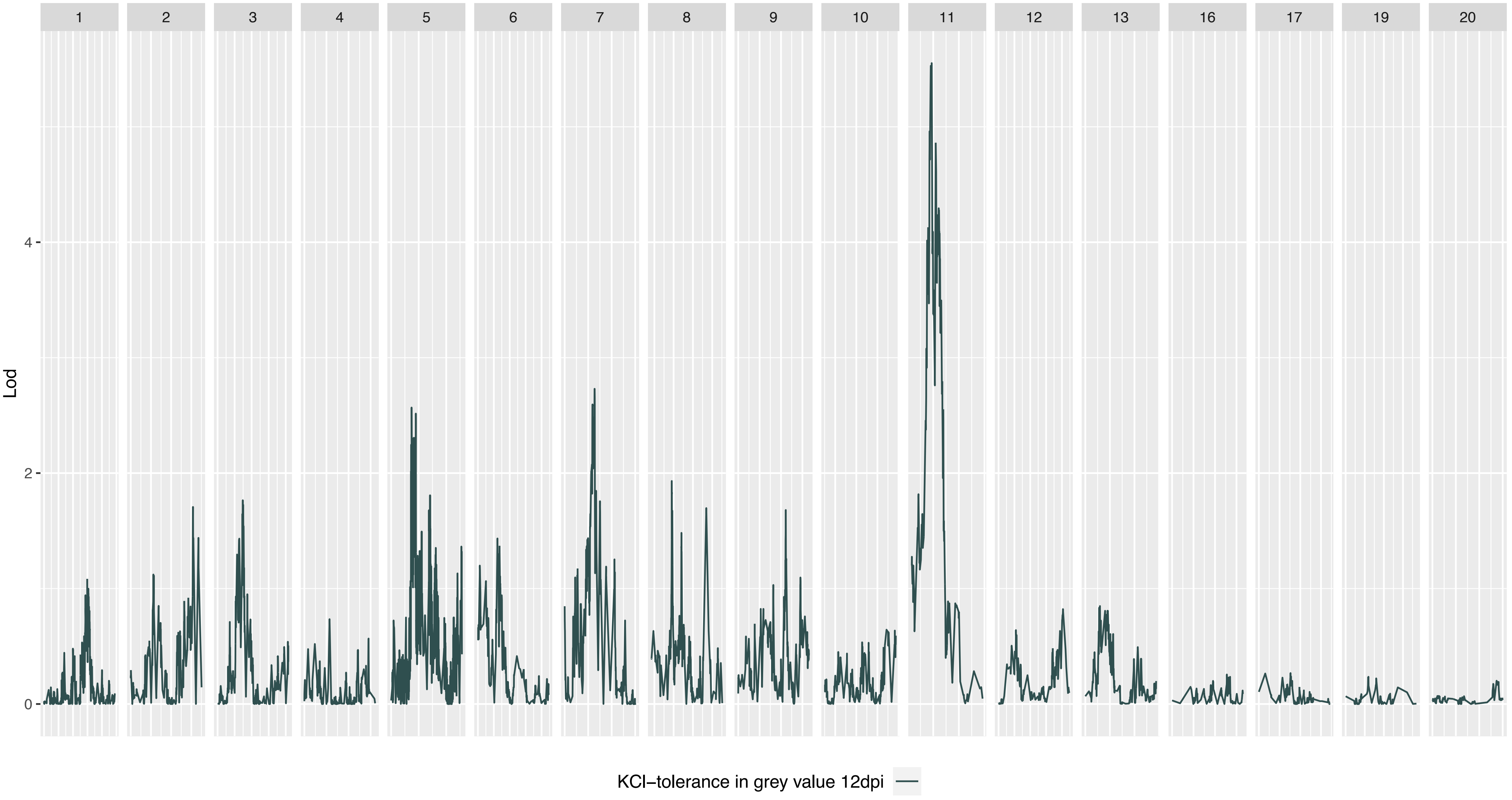

1
2
3
4
5
6
7
8
9
10
11
12
13
16
17
19
20
Lod
4
2
0
KCl−tolerance in grey value 12dpi

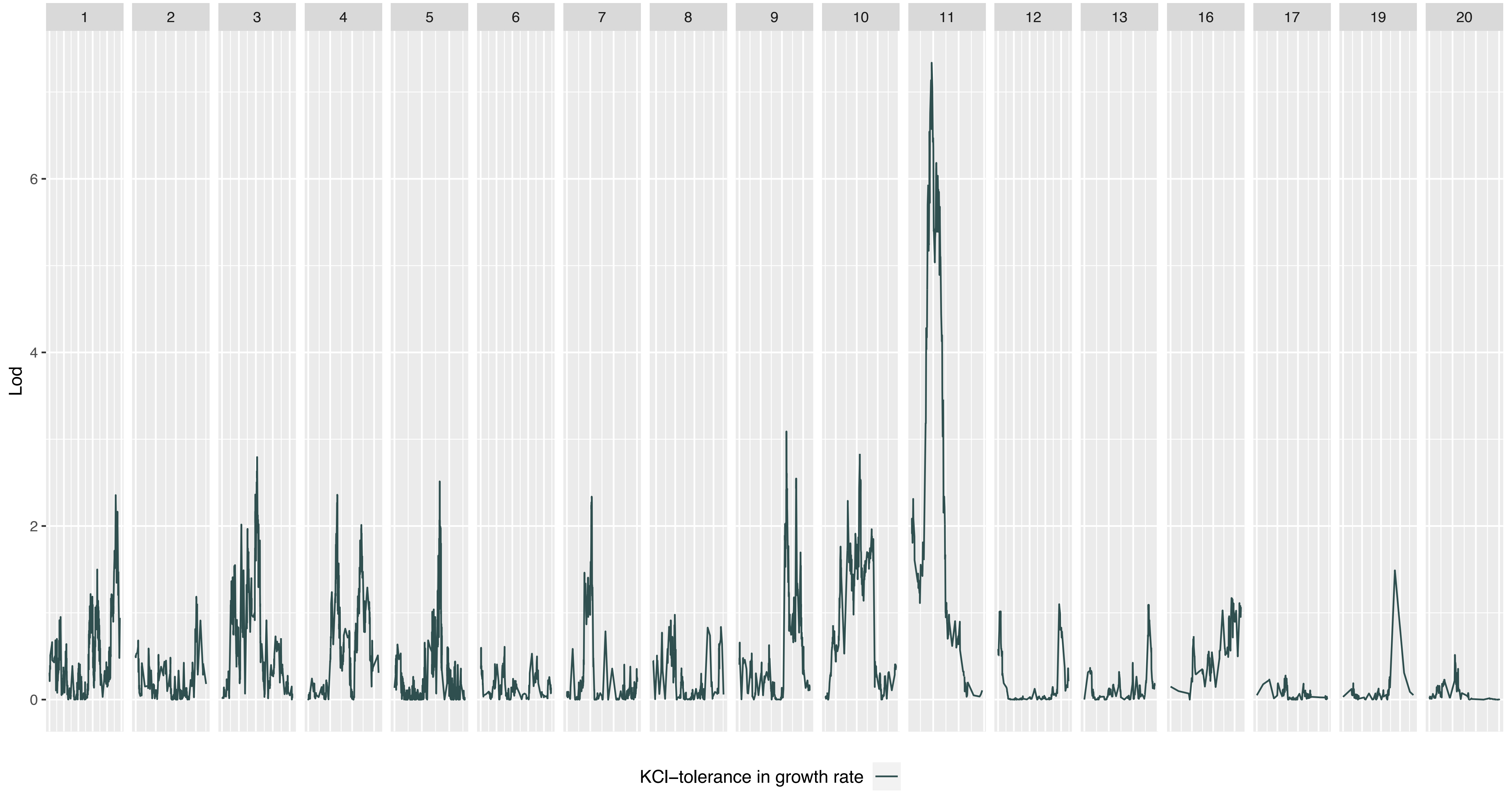
1
2
3
4
5
6
7
8
9
10
11
12
13
16
17
19
20
Lod
0
2
4
6
KCl–tolerance in growth rate

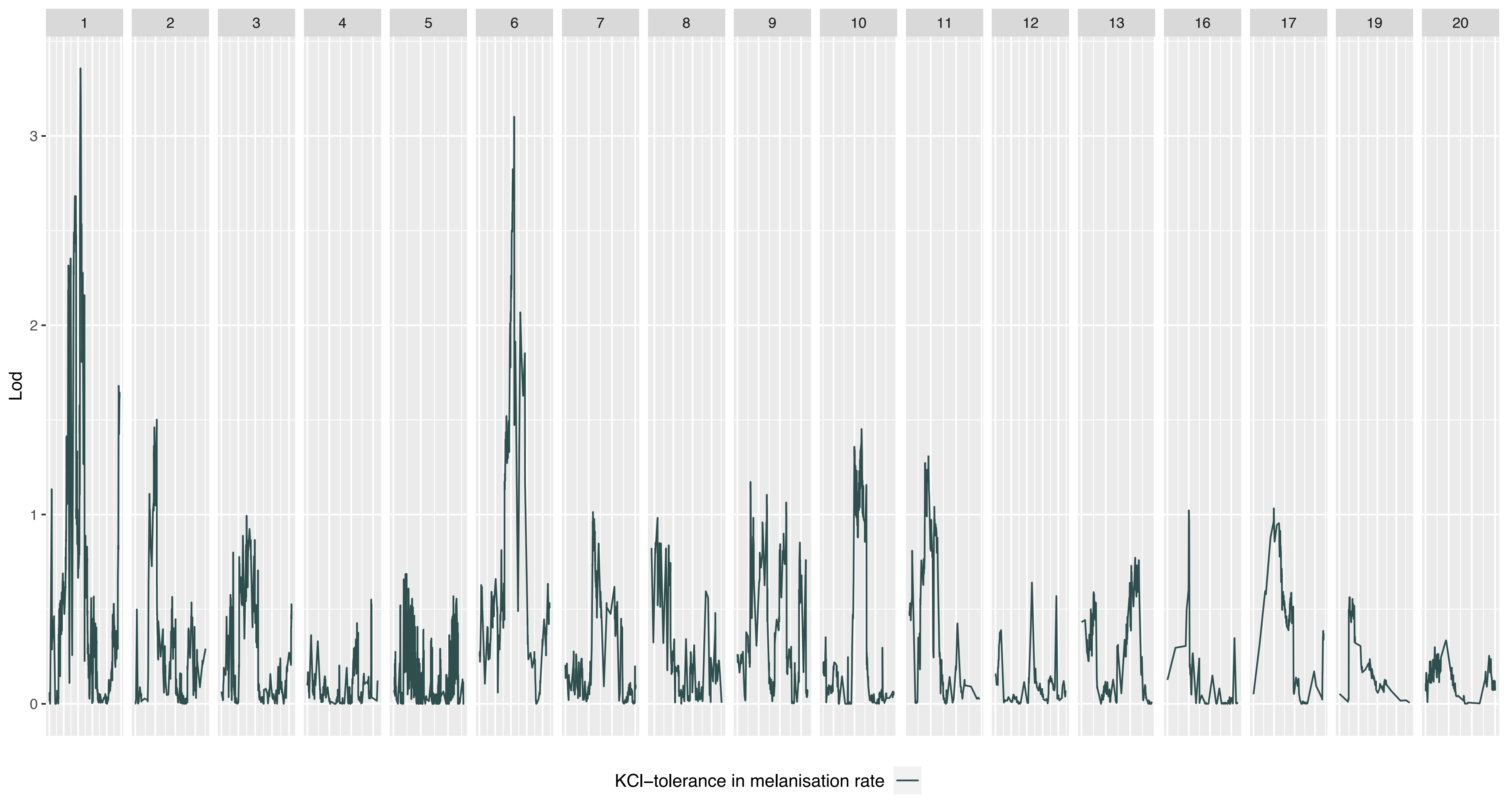
1
2
3
4
5
6
7
8
9
10
11
12
13
16
17
19
20
Lod
3
2
1
0
KCl−tolerance in melanisation rate
