## Supplementary material for "Quantitative trait locus mapping of osmotic stress response in the fungal wheat pathogen *Zymoseptoria tritici*": Figure S4

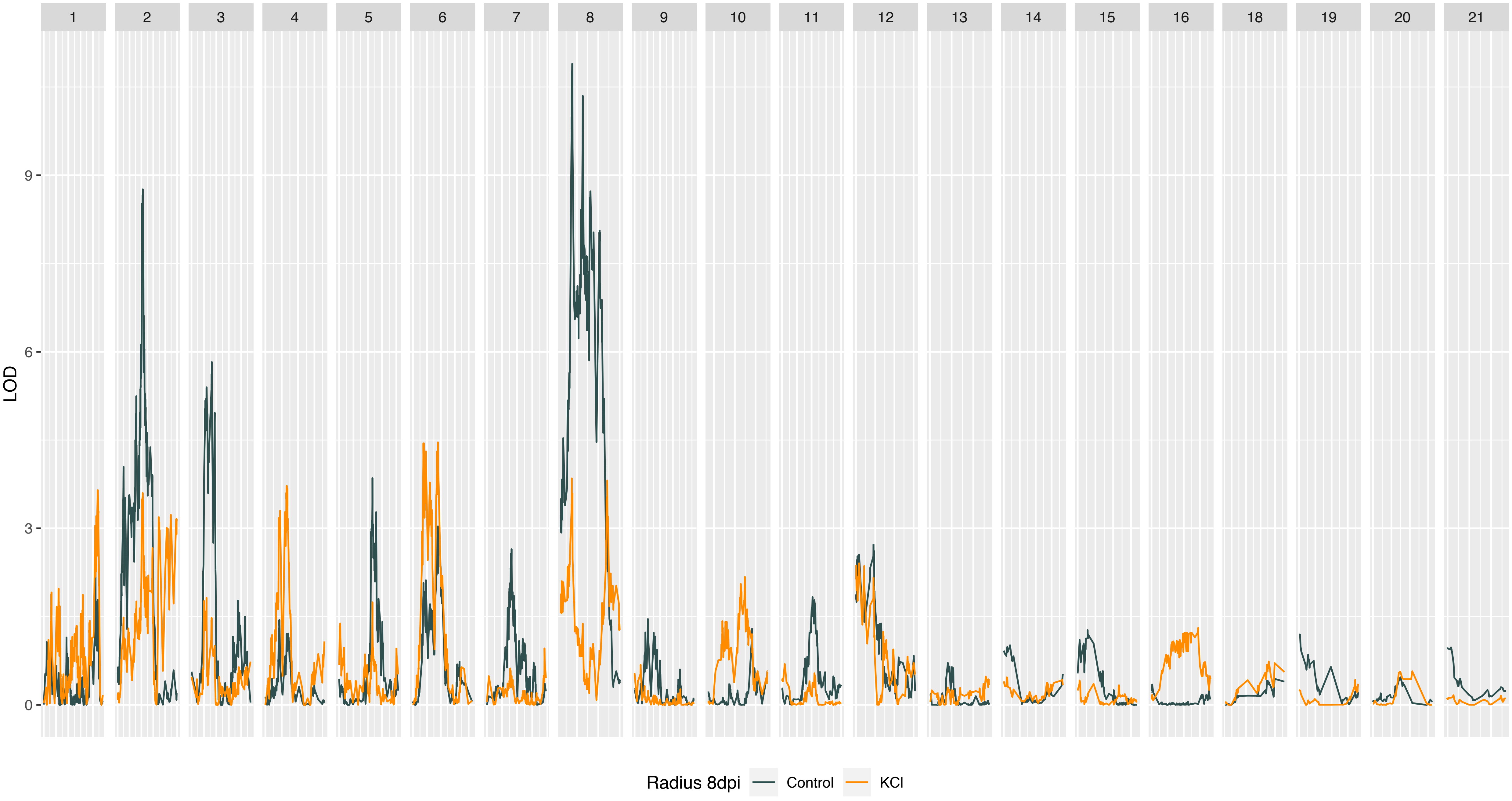

18
19
20
21
LOD
0
3
6
9
Radius 8dpi
Control
KCl

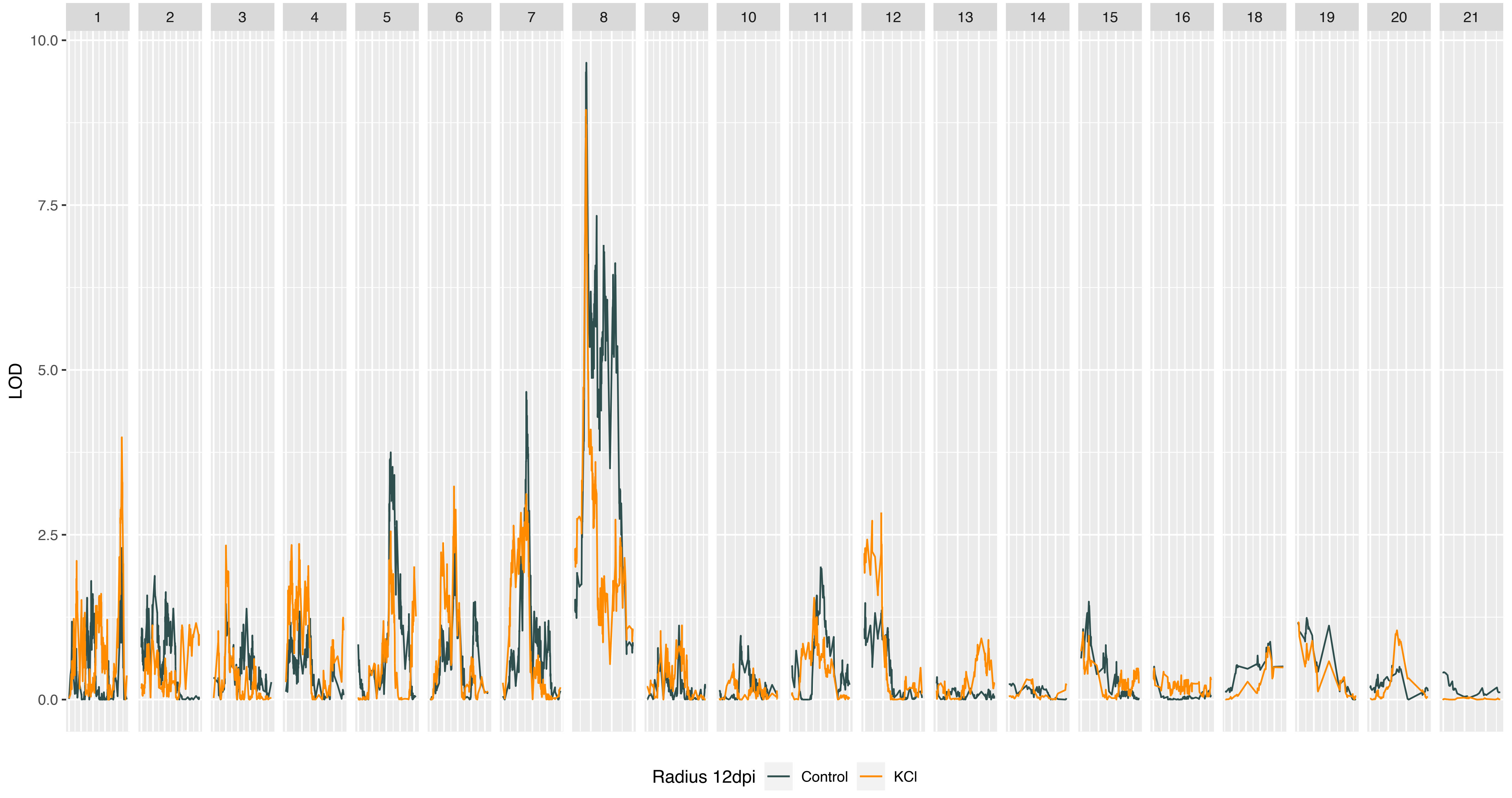
18
19
20
21
LOD
10.0
7.5
5.0
2.5
0.0
Radius 12dpi
Control
KCl

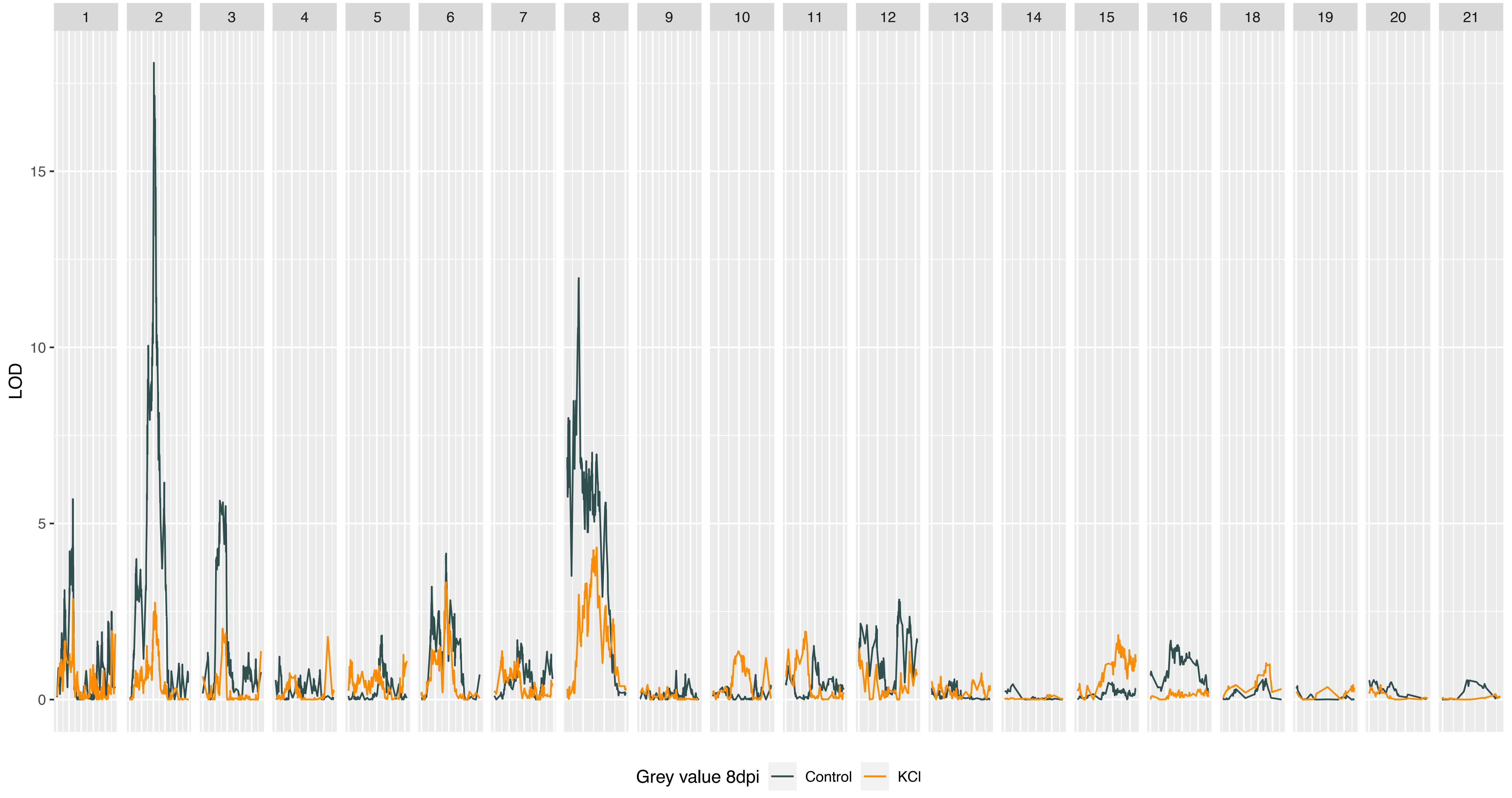
18
19
20
21
LOD
15
10
5
0
Grey value 8dpi
Control
KCl

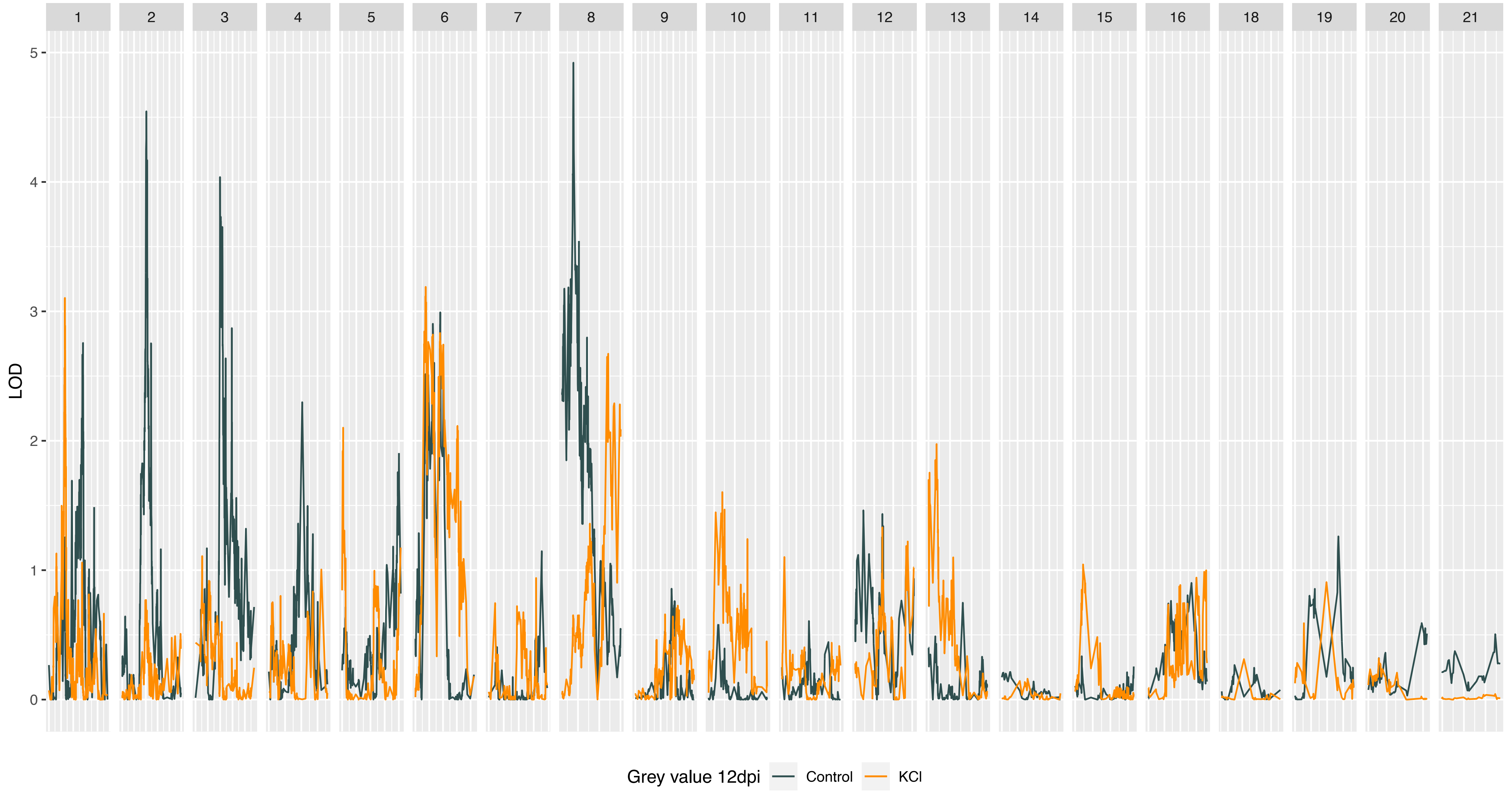

18
19
20
21
LOD
5
4
3
2
1
0
Grey value 12dpi
Control
KCl

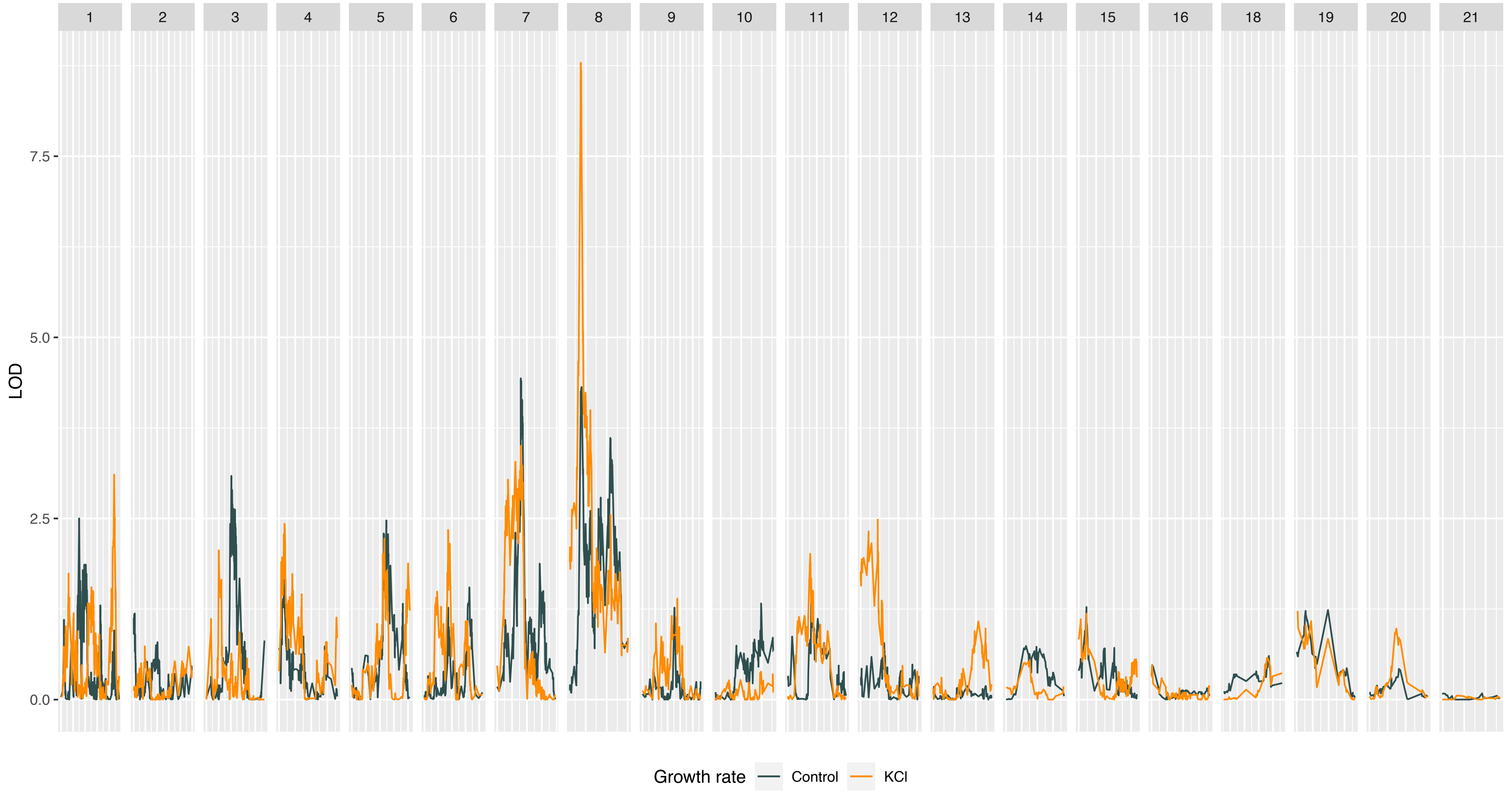
18
19
20
21
LOD
7.5
5.0
2.5
0.0
Growth rate
Control
KCl

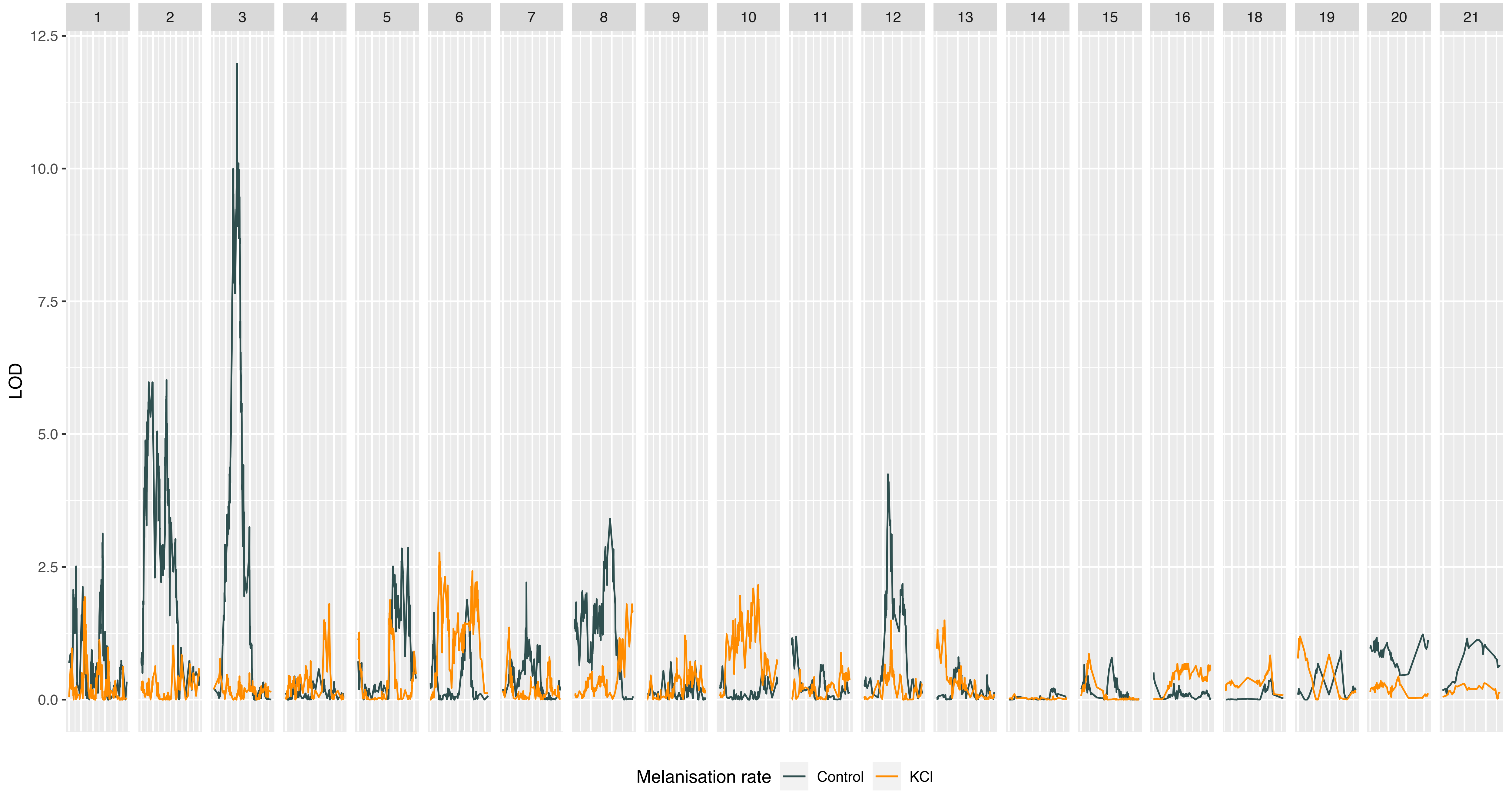
18
19
20
21
LOD
12.5
10.0
7.5
5.0
2.5
0.0
Melanisation rate
Control
KCl

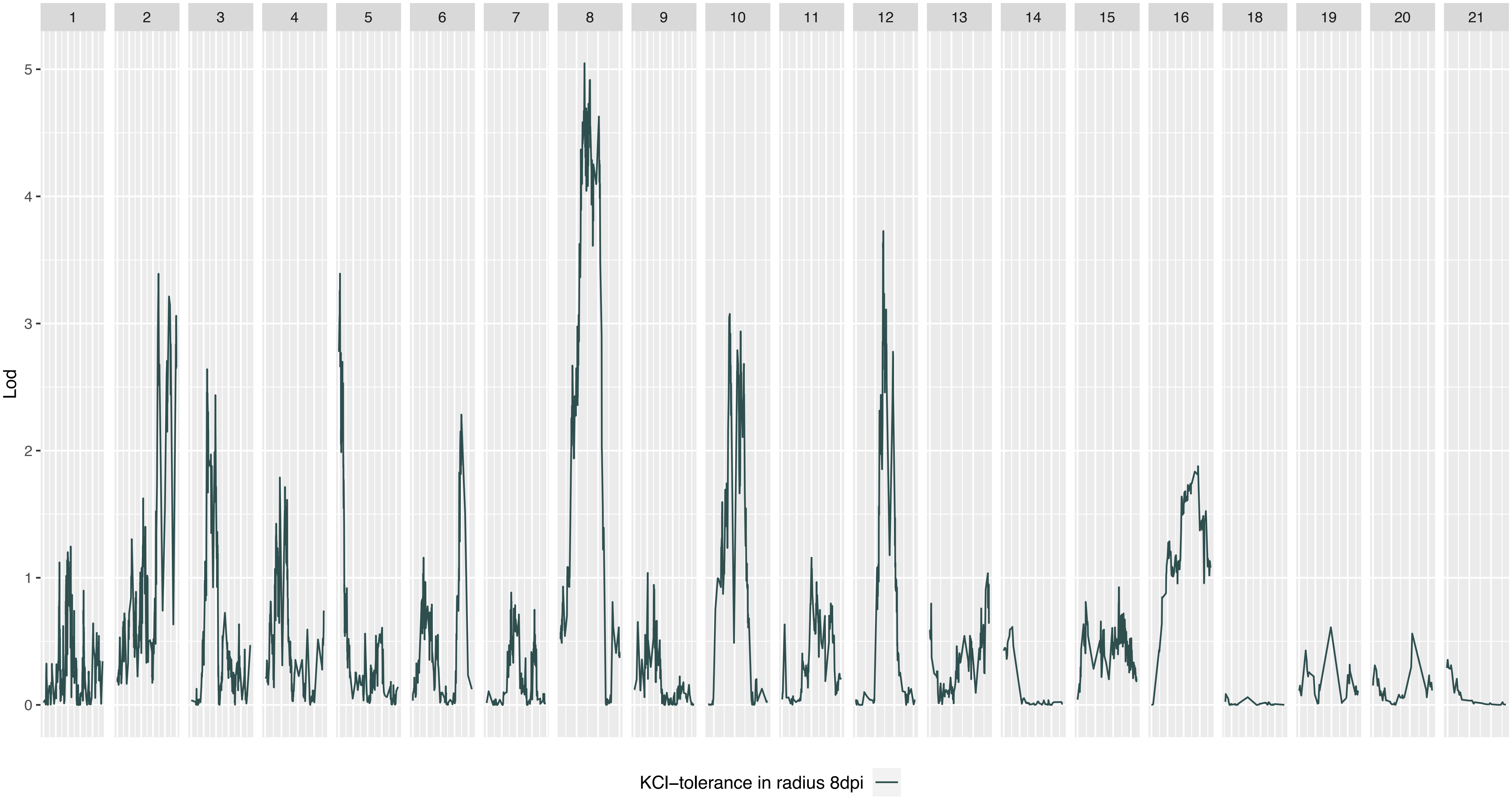

18
19
20
21
Lod
0
1
2
3
4
5
KCl–tolerance in radius 8dpi

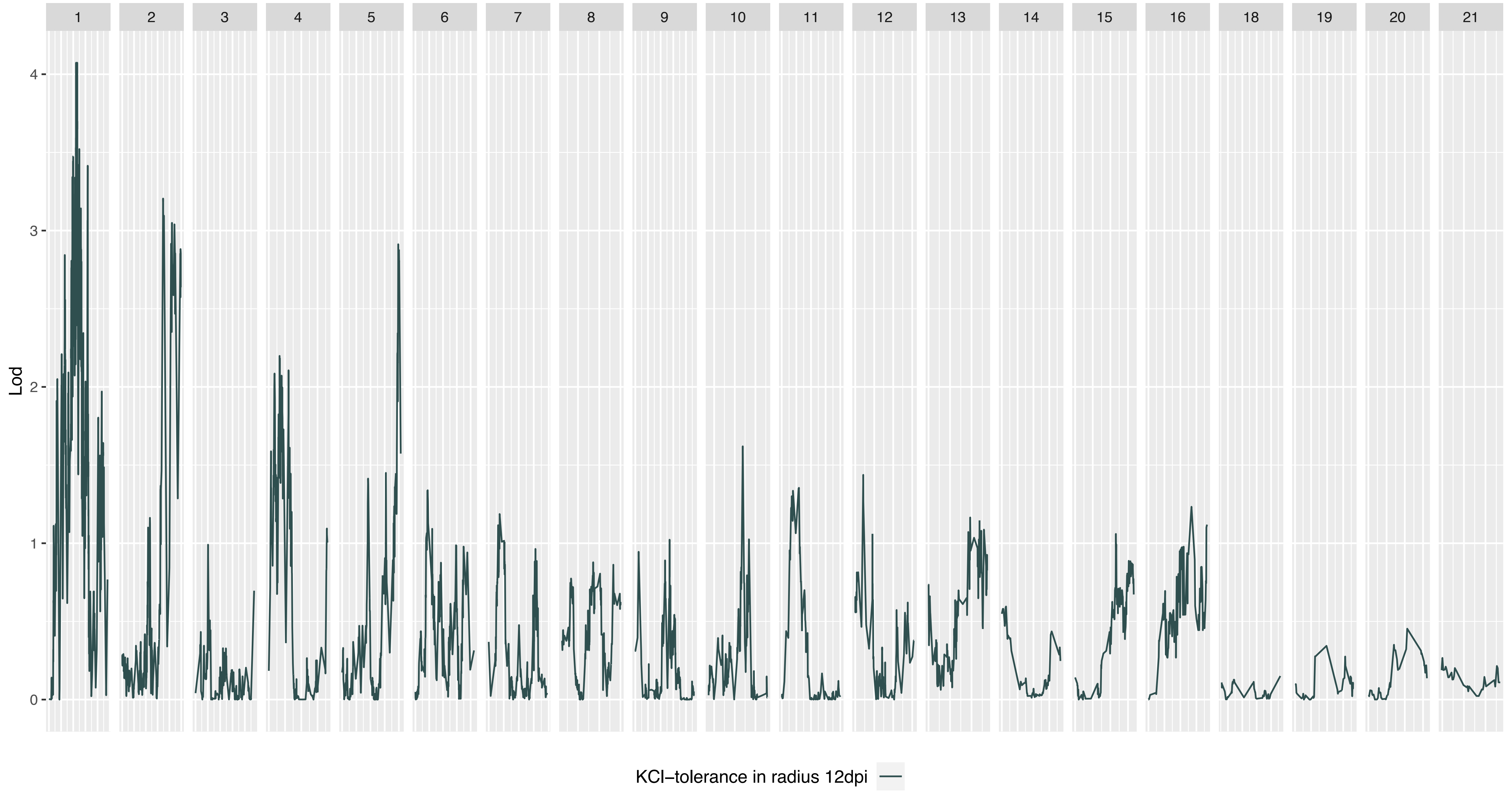
18
19
20
21
Lod
0
1
2
3
4
KCl–tolerance in radius 12dpi

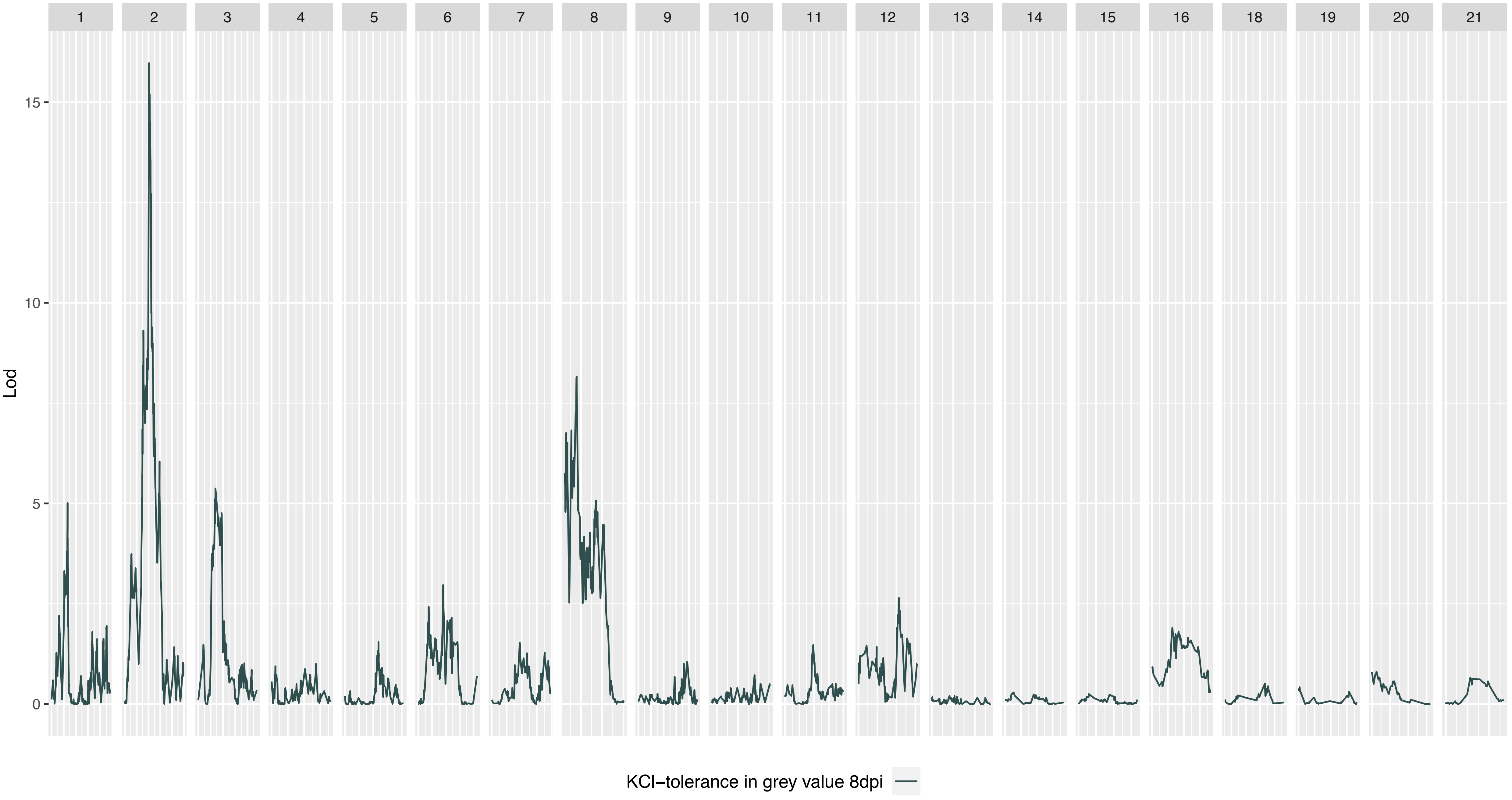

18
19
20
21
Lod
0
5
10
15
KCl−tolerance in grey value 8dpi

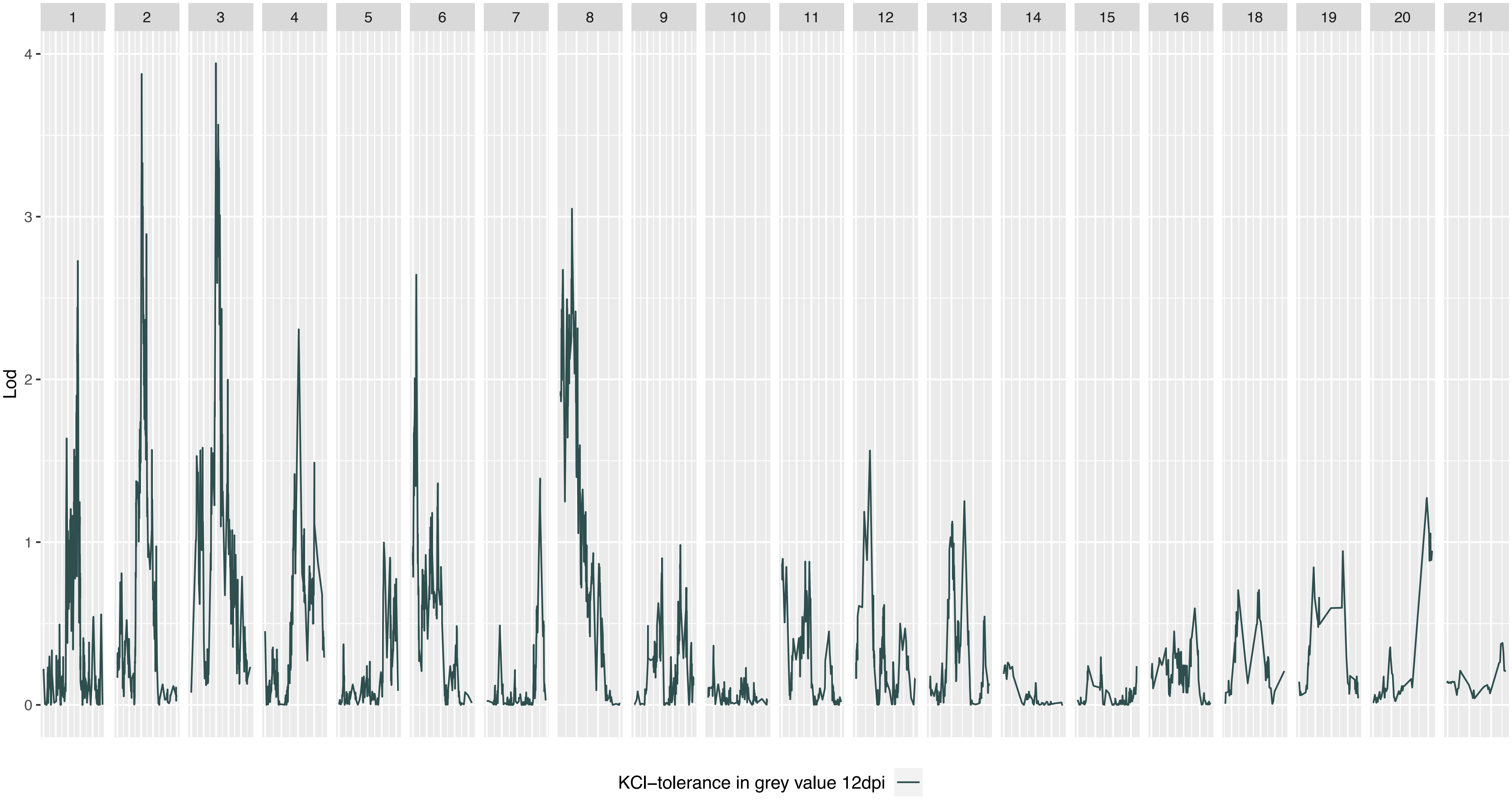
18
19
20
21
Lod
0
1
2
3
4
KCl–tolerance in grey value 12dpi

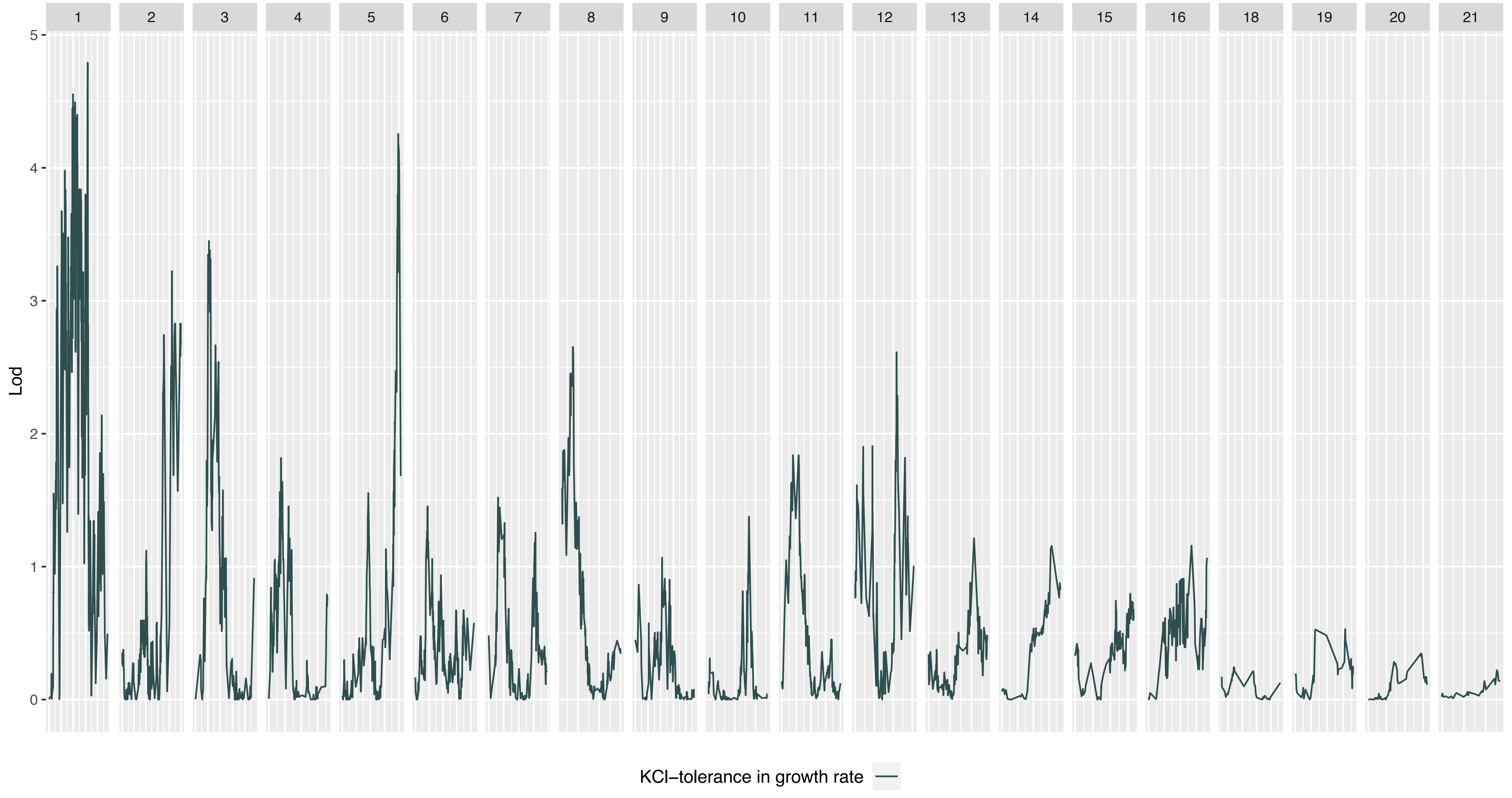
18
19
20
21
Lod
5
4
3
2
1
0
KCl–tolerance in growth rate

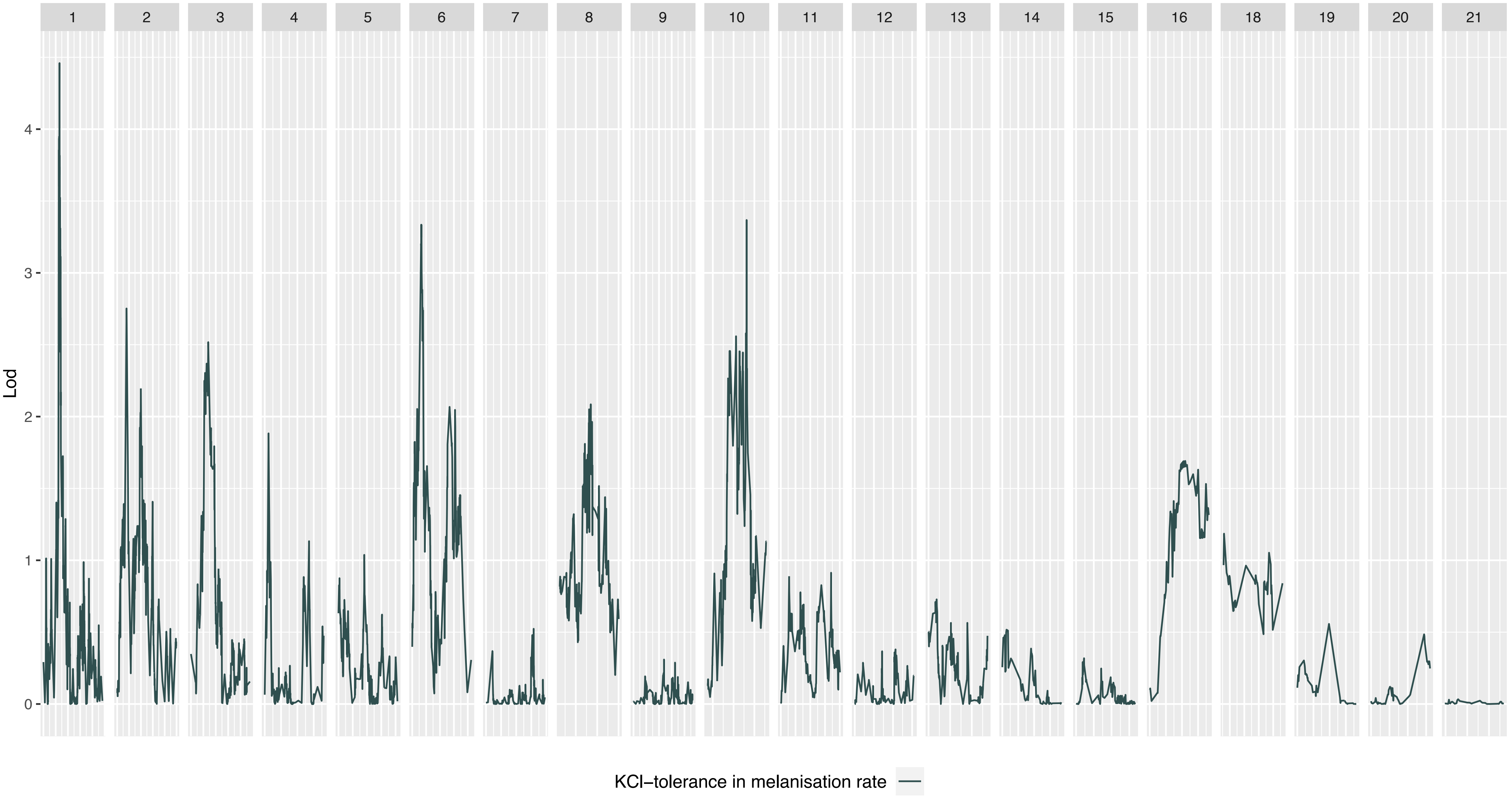
18
19
20
21
Lod
4
3
2
1
0
KCl–tolerance in melanisation rate
